# *Prevotella copri*-related effects of a therapeutic food for malnutrition

**DOI:** 10.1101/2023.08.11.553030

**Authors:** Hao-Wei Chang, Evan M. Lee, Yi Wang, Cyrus Zhou, Kali M. Pruss, Suzanne Henrissat, Robert Y. Chen, Clara Kao, Matthew C. Hibberd, Hannah M. Lynn, Daniel M. Webber, Marie Crane, Jiye Cheng, Dmitry A. Rodionov, Aleksandr A. Arzamasov, Juan J. Castillo, Garret Couture, Ye Chen, Nikita P. Balcazo, Carlito B. Lebrilla, Nicolas Terrapon, Bernard Henrissat, Olga Ilkayeva, Michael J. Muehlbauer, Christopher B. Newgard, Ishita Mostafa, Subhasish Das, Mustafa Mahfuz, Andrei L. Osterman, Michael J. Barratt, Tahmeed Ahmed, Jeffrey I. Gordon

## Abstract

Preclinical and clinical studies are providing evidence that the healthy growth of infants and children reflects, in part, healthy development of their gut microbiomes^1–5^. This process of microbial community assembly and functional maturation is perturbed in children with acute malnutrition. Gnotobiotic animals, colonized with microbial communities from children with severe and moderate acute malnutrition, have been used to develop microbiome-directed complementary food (MDCF) formulations for repairing the microbiomes of these children during the weaning period^5^. Bangladeshi children with moderate acute malnutrition (MAM) participating in a previously reported 3-month-long randomized controlled clinical study of one such formulation, MDCF-2, exhibited significantly improved weight gain compared to a commonly used nutritional intervention despite the lower caloric density of the MDCF^6^. Characterizing the ‘metagenome assembled genomes’ (MAGs) of bacterial strains present in the microbiomes of study participants revealed a significant correlation between accelerated ponderal growth and the expression by two *Prevotella copri* MAGs of metabolic pathways involved in processing of MDCF-2 glycans^1^. To provide a direct test of these relationships, we have now performed ‘reverse translation’ experiments using a gnotobiotic mouse model of mother-to-offspring microbiome transmission. Mice were colonized with defined consortia of age- and ponderal growth-associated gut bacterial strains cultured from Bangladeshi infants/children in the study population, with or without *P. copri* isolates resembling the MAGs. By combining analyses of microbial community assembly, gene expression and processing of glycan constituents of MDCF-2 with single nucleus RNA-Seq and mass spectrometric analyses of the intestine, we establish a principal role for *P. copri* in mediating metabolism of MDCF-2 glycans, characterize its interactions with other consortium members including *Bifidobacterium longum* subsp. *infantis*, and demonstrate the effects of *P. copri-*containing consortia in mediating weight gain and modulating the activities of metabolic pathways involved in lipid, amino acid, carbohydrate plus other facets of energy metabolism within epithelial cells positioned at different locations in intestinal crypts and villi. Together, the results provide insights into structure/function relationships between MDCF-2 and members of the gut communities of malnourished children; they also have implications for developing future prebiotic, probiotic and/or synbiotic therapeutics for microbiome restoration in children with already manifest malnutrition, or who are at risk for this pervasive health challenge.

---

In a recent study^1^, we describe results obtained from a metagenomic analysis of fecal samples serially collected from 12-18-month-old Bangladeshi children with moderate acute malnutrition enrolled in a randomized clinical trial that tested the effects of administering a microbiota-directed complementary food prototype (MDCF-2) versus a ready-to-use supplementary food (RUSF) on host physiology and the structure and expressed functions of their microbiomes. Sequencing reads from fecal DNA were assembled into bacterial MAGs and MAGs were identified whose abundances were associated with anthropometric measures of ponderal growth (changes in weight-for-length (height), expressed as WLZ scores). Changes in MAG gene expression were also characterized in response to treatment and related to the metabolism of carbohydrate components of MDCF-2. A notable finding was that among the 75 MAGs found to be positively correlated with WLZ, two belonging to *Prevotella copri* were a principal source of expression of carbohydrate utilization pathways that target MDCF-2 polysaccharides. Moreover, expression of these pathways and the levels of products of glycan metabolism were significantly positively correlated with the magnitude of the childrens’ improvement in WLZ^1^.

Here we describe ‘reverse translation’ experiments, conducted in gnotobiotic mice, that further examine the role of *P. copri* and other community members in mediating MDCF-2 metabolism, and the effects of this therapeutic food on aspects of host physiology, notably weight gain and intestinal function. These experiments involved sequential introduction, during the suckling and weaning periods of defined collections of genome-sequenced bacterial strains, cultured from the study population, into gnotobiotic female mice (dams)with subsequent transmission of these strains to their pups. The strains represented organisms whose prominence normally changes at different stages of postnatal gut microbial community assembly in healthy Bangladeshi children (‘age-discriminatory’ taxa)^4,5,8,9^; they also include WLZ-correlated bacterial strains selected based on their shared features with MAGs identified in the clinical study^1,6^. Pups were subjected to the same dietary sequence of exclusive milk feeding (from the dam) followed by weaning onto an MDCF-2 supplemented diet.

The small intestine is lined with a layer of continuously renewing epithelial cells. Renewal is fueled by Lgr5^+^ stem cells located at the base of mucosal invaginations (crypts of Lieberkühn). Stem cell progenies adopt four principal fates. Members of the predominant enterocytic lineage execute their differentiation program as they migrate from crypts up adjacent small intestinal villi. Goblet cells, enteroendocrine cells, tuft cells, and Paneth cells represent other lineal descents of the multipotential crypt stem cell and its oligo-potential daughters. Substantial metabolic investments are needed to support the normal daily replacement of large numbers of gut epithelial cells^7^ as well as the functions they normally express in their differentiating and differentiated states. Therefore, we tested the hypothesis that epithelial cell gene expression and metabolism would be sensitive reporters of functional differences associated with defined consortia that did or did not contain *P. copri* strains representing the MAGs. The results highlight a central role played by *P. copri* strains closely resembling the identified WLZ-associated MAGs in metabolizing glycans present in MDCF-2, plus their capacity, in the context of this consortium of strains and MDCF-2, to promote weight gain and influence expression of metabolic functions and other activities in enterocytes as they migrate up the villus.

## Results

### A manipulatable model of mother-pup transmission of cultured age- and WLZ-associated bacterial taxa

#### Selection of bacterial strains

To test the role of *P. copri* in the context of a defined human gut microbial community that captured features of the developing communities of children who had been enrolled in the clinical study of MDCF-2^1^, we selected 20 bacterial strains, 16 of which were cultured from the fecal microbiota of 6- to 24-month-old Bangladeshi children living in Mirpur, the urban slum where the previously reported randomized controlled MDCF-2 clinical trial had been performed (**Supplementary Table 1a**). They included strains initially identified by the close correspondence of their 16S rRNA gene sequences to (i) a group of taxa that describe a normal program of development of the microbiota in healthy Bangladeshi children^8,9^ and (ii) taxa whose abundances had statistically significant associations (positive or negative) with the rate of weight gain (β-WLZ) and statistically significant correlations with plasma levels of WLZ-associated proteins in children from the clinical study^5,6^. The relatedness of these strains to the 1,000 MAGs assembled from fecal samples obtained from all participants in the clinical study^1^ was determined by average nucleotide sequence identity (ANI) scores, alignment coverage parameters^10,11^ and their encoded metabolic pathways (**Supplementary Table 1b-d**). Encoded metabolic pathways for carbohydrate utilization, amino acid and vitamin/cofactor biosynthesis and fermentation represented in the MAGs and cultured strains, were defined by *in silico* reconstructions and the results described in the form of ‘binary phenotype’ scores denoting pathway presence or absence^1^.

Liquid chromatography-mass spectrometry analysis of glycosidic linkages and polysaccharides in MDCF-2 and RUSF disclosed that cellulose, galactan, arabinan, xylan, and mannan represent the principal non-starch polysaccharides in MDCF-2. Fecal microbial RNA-seq datasets generated from clinical study participants revealed that in comparisons of (i) MDCF-2 vs. RUSF treatment and (ii) children who had received MDCF-2 and exhibited the greatest vs. lowest (upper vs. lower quartile) WLZ responses, there was significant enrichment of expression of genes involved in utilization of MDCF-2 glycans, including arabinose and α-arabinooligosaccharides (aAOS). Two *P. copri* MAGs (Bg0018 and Bg0019) whose abundances were positively correlated with WLZ contributed a large share of the genes driving this enrichment. Comparisons of the polysaccharide utilization loci (PULs) represented in the 11 *P. copri* MAGs detected in the study population revealed that Bg0018 and Bg0019 share 10 functionally conserved (PULs), including seven that were completely conserved (see *Methods* and ref. 1 for the criteria used to classify the degree of PUL conservation). These 10 PULs encode a diverse set of glycoside hydrolases with broad substrate specificities for glycans present in MDCF-2 (**Supplementary Table 1e**). Notably, the degree of representation of the seven completely conserved PULs among the 11 *P. copri* MAGs identified in study participants was predictive of each MAG’s strength of association with WLZ, suggesting a link between metabolism of carbohydrates by *P. copri* and ponderal growth responses among these children^1^.

To test how *P. copri* colonization with a strain resembling MAGs Bg0018 and Bg0019 affected microbial community composition and expressed functions, dietary glycan degradation, and host metabolism, we selected the Bangladeshi *P. copri* strain PS131.S11 (abbreviated *P. copri* Bg131). This strain was chosen over four other isolates cultured from Bangladeshi children due to its phylogenetic similarity to Bg0018 and Bg0019 (**Extended Data Fig. 1a**), concordance of its metabolic pathway representation with these MAGs (**Supplementary Table 1c,d**) and its representation of five of the 10 functionally conserved PULs shared by Bg0018 and Bg0019. These five PULs are predicted to be involved in degradation of starch, ß-glucan, pectin, pectic galactan, and xylan (**Supplementary Table 1e,f)**. An additional PUL targeting arabinogalactan was found adjacent to the conserved PUL targeting starch, though it did not meet criteria for conservation with the corresponding PUL in MAGs Bg0018 and Bg0019 (**Supplementary Table 1f**).

To assess the specificity of responses of *P. copri* to MDCF-2, we additionally included an isolate from another *Prevotella* species, *P. stercorea*. *P. stercorea* did not have any WLZ-associated MAGs identified in the clinical study and the cultured isolate did not share any of the PULs present in MAGs Bg0018 and Bg0019 or *P. copri* Bg131. Instead, the PULs in the *P. stercorea* isolate contain glycoside hydrolases mainly targeting animal-derived glycans (**Supplementary Table 1g**). Therefore, we hypothesized this isolate would exhibit lower fitness on a plant glycan-based diet of MDCF-2.

*Bifidobacterium longum* subsp. *infantis* (*B. infantis*) is a prominent early colonizer of the infant gut. Therefore, we wanted to ensure that it was well represented at the earliest stages of assembly of the defined community so that later colonizers such as *P. copri* could establish themselves. The collection of cultured isolates included two strains of *B. infantis* recovered from Bangladeshi children – *B. infantis* Bg463 and *B. infantis* Bg2D9. The Bg463 strain had been used in our earlier preclinical studies that led to development of MDCF-2^5,9^. In a recent study, simultaneous introduction of five different Bangladeshi *B. infantis* strains, without any other organisms, into just-weaned germ-free mice fed a diet representative of that consumed by 6-month-old children living in Mirpur^12^ revealed that *B. infantis* Bg2D9 exhibited greater fitness (absolute abundance) than any of the other strains, including Bg463. Based on comparative genomic and microbial RNA-Seq analyses, this improved fitness was attributed to additional carbohydrate utilization pathways that the Bg2D9 strain possesses^12^. In addition, a combination of *B. infantis* Bg2D9 and a commercial *B. infantis* strain was found to promote weight gain when introduced into mice colonized with the intact uncultured fecal microbiota of an infant with severe acute malnutrition^12^.

### Establishing *P. copri* colonization and assessing effects of different *B. infantis* strains

#### Design

We used the 20-strain collection to perform a 3-arm, fixed diet study that involved ‘successive’ waves of maternal colonization with four different bacterial consortia (**Fig. 1a-c**). The sequence of introduction of taxa into dams was designed to emulate temporal features of the normal postnatal development of the human gut community; *e.g*., consortia 1 and 2 were comprised of strains that are prominent colonizers of healthy infants/children in the first postnatal year (including the *B. infantis* isolates) while those in consortium 3 are prominent during weaning in the second postnatal year^4,5,8,9^. This dam-to-pup colonization strategy also helped overcome the technical challenge of reliable delivery of bacterial consortia to newborn pups via oral gavage.

**Fig. 1.**
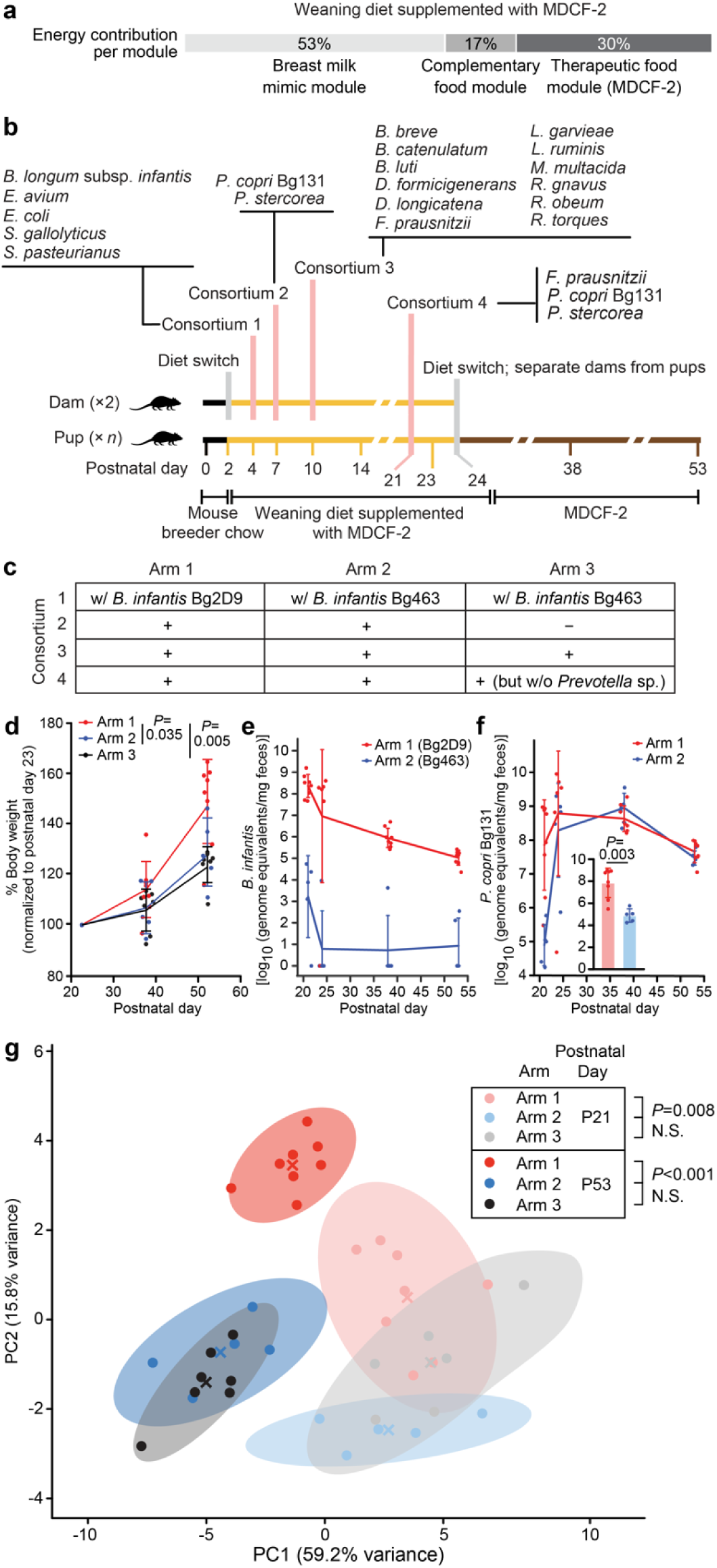
Identifying factors that affect the efficiency of colonization of gnotobiotic dam-pup dyads with *P. copri* in the presence of other cultured age-discriminatory and WLZ-associated bacterial strains and the effects of colonization on pup weight gain. **(a)** Energy contribution from different modules of the ‘weaning diet supplemented with MDCF-2’. (**b,c**) Study design (n=2 dams and 5-8 offspring/treatment arm). Panel b outlines the timing of bacterial colonization of dams and diet switches. Panel c describes the gavages administered to members of each treatment arm. (**d**) Body weights of the offspring of dams, normalized to postnatal day 23. [linear mixed effects model (see *Methods*)]. Mean values ± SD are shown. Each dot in panels d-f represents an individual animal. *P*-values were calculated using a Mann-Whitney U test (panel e inset) or a linear mixed effect model (panel f). (**e**) Absolute abundance of *B. infantis* Bg2D9 (Arm 1) and *B. infantis* Bg463 (Arm 2) in fecal samples obtained from pups. (**f**) Absolute abundance of *P. copri* in fecal samples collected from pups in the indicated treatment arms at the indicated postnatal time points. Inset: the absolute abundance of *P. copri* in fecal samples collected from pups at P21. (**g**) Principal components analysis of absolute abundances of other community members in fecal samples obtained from pups at postnatal day 21 and postnatal day 53. Centroids are denoted by a colored “X”. Shaded ellipses represent the 95% confidence interval of the sample distribution. *P*-values were calculated with PERMANOVA on PC projections. Each dot represents an individual animal.

Dually-housed germ-free dams were switched from a standard breeder chow to a ‘weaning-diet’ supplemented with MDCF-2 on postpartum day 2, two days before initiation of the colonization sequence. This weaning diet was formulated to emulate the diets consumed by children in the clinical trial during MDCF-2 treatment (See *Methods*; **Fig 1a; Supplementary Table 2**); it contained (i) powdered human infant formula, (ii) complementary foods consumed by 18-month-old children living in Mirpur, Bangladesh where the study took place, and (iii) MDCF-2. The contributions of the milk, complementary food and MDCF-2 ‘modules’ to total caloric content (53%, 17%, and 30%, respectively) were based on published studies of the diets of cohorts of healthy and undernourished 12- to 23-month-old children from several low- and middle-income countries, including Bangladesh, as well as the amount of MDCF-2 given to the 12-18-month-old children with MAM in the clinical study^1,5,6,13–16^. Pups in all three arms were subjected to a diet sequence that began with exclusive milk feeding (from the nursing dam) followed by a weaning period where pups had access to the weaning phase diet supplemented with MDCF-2. Pups were weaned at postnatal day 24 (P24), after which time they received MDCF-2 alone *ad libitum* until P53 when they were euthanized.

Initial attempts to mono-colonize mice with *P. copri* revealed that it is a poor colonizer on its own (**Extended Data Fig. 1b,c**). To identify factors that promote stable *P. copri* colonization, we performed an experiment comparing the effects of the two different *B. infantis* strains, Bg2D9 and Bg463, on *P. copri* and the other cultured strains representing age- and WLZ-associated MAGs. We included *B. infantis* Bg2D9 in consortium 1 of Arm 1 of the 3-arm experiment and *B. infantis* Bg463 in consortium 1 of Arm 2 (**Fig. 1b,c**). This consortium consisted of five ‘early’ infant gut community colonizers and was administered to dams on postpartum day 4. Dams in these two arms subsequently received the following gavages: (i) on postpartum day 7, *P. copri* and *P. stercorea*; (ii) on postpartum day 10, additional age-discriminatory and WLZ-associated taxa, and (iii) on postpartum day 21, three strains - *P. copri, P. stercorea,* and *Faecalibacterium prausnitzii* (**Fig. 1b**). At this last time point, the three strains were given by oral gavage to both the dams and their offspring to help promote successful colonization. To test the effects of *P. copri* colonization, we included Arm 3, which was a replicate of Arm 2 but without *Prevotella* in consortia 2 and 4. Our rationale for the timing of the first three gavages was based on the diet sequence [gavage 1 of early colonizers at a time (P4) when mice were exclusively consuming the dam’s milk, gavage 2 as the pups were just beginning to consume the human weaning (complementary food) diet, gavage 3 somewhat later during this period of ‘complementary feeding and the fourth gavage to help to ensure a consistent level of *P. copri* colonization at the end of weaning (and subsequently through the post-weaning period)].

#### Effects of B. infantis on weight gain, P. copri colonization, and community composition

Gnotobiotic mice colonized with *B. infantis* Bg2D9 exhibited a significantly greater increase in weight gain between P23 (the first time point measured, 2 days after the final gavage) and P53 compared to mice in the two other experimental arms, supporting prior observations of its ability to promote ponderal growth^12^ [*P* < 0.05 for Arm 1 versus Arm 2; *P*<0.01 for Arm 1 versus Arm 3; linear mixed-effects model (see *Methods*)] (**Fig. 1d**). In contrast, there was no significant difference in weight gain between animals colonized with *B. infantis* Bg463 with or without *Prevotella* species in Arms 2 and 3.

To explore whether these differences in weight gain were associated with differences in microbial community composition, we quantified the relative and absolute abundances of all administered strains in fecal samples collected from dams on days postpartum days 21, 24, and 35, fecal samples collected from their offspring on P21, P24, P35, and P53, as well as cecal and ileal contents collected from offspring at P53 by shotgun sequencing of community DNA (n=2 dams and 5-8 pups analyzed/arm; **Supplementary Table 3a**). Colonization of each consortium member was highly consistent among all animals in each treatment group (**Supplementary Table 3b-d)**. *B. infantis* Bg2D9 successfully colonized pups at P21 in Arm 1 [8.4±0.5 log_10_ (genome equivalents/mg feces) (mean±SD); relative abundance, 9.0±3.9% (mean±SD)]. In contrast, *B. infantis* Bg463 colonized at 5-8 orders-of-magnitude lower absolute abundance levels in Arms 2 and 3 [3.22±1.9 and 0.6±1.5 log_10_ (genome equivalents/mg feces) (mean±SD), respectively]. These differences between the groups were sustained through P53. Consistent with the role of *B. infantis* as an early pre-weaning colonizer, both strains decreased in abundance between P21 and P53 (**Fig. 1e**). Exposure to *B. infantis* Bg2D9 in Arm 1 was associated with levels of *P. copri* colonization that were 3-orders of magnitude greater in the pre-weaning period (P21) than in Arm 2 mice exposed to *B. infantis* Bg463 [*P*<0.005, Mann-Whitney U test] (**Fig. 1f; Supplementary Table 3c**). Administering the fourth gavage on P21 elevated the absolute abundance of fecal *P. copri* in Arm 2 to a level comparable to Arm 1; this level was sustained throughout the post-weaning period (P24 to P53) [**Fig. 1f**; *P*>0.05; linear mixed effects model (*Methods*); also see **Supplementary Table 3c**]. This effect of the fourth gavage was also evident in the ileal and cecal microbiota at the time of euthanasia (**Supplementary Table 3d**).

Based on these results, we directly tested the colonization dependency of *P. copri* on *B. infantis*, in two independent experiments whose designs are outlined in **Extended Data Fig. 1b**. Dually-housed germ-free dams were switched from standard breeder chow to the weaning Bangladeshi diet supplemented with MDCF-2 on postpartum day 2. On postpartum day 4, one group of dams was colonized with *B. infantis* Bg2D9 while the other group received a sham gavage. On postpartum days 7 and 10, both groups of gnotobiotic mice were gavaged with a consortium containing five *P. copri* strains. These five *P. copri* strains (Bg131, 1A8, 2C6, 2D7 and G8) were all isolated from fecal samples obtained from Bangladeshi children (**Supplementary Table 4**). Pups were separated from their dams at the completion of weaning and their diet was switched to MDCF-2 until euthanasia on P42 (n=9 mice per treatment group; 2 independent control experiments). At this time point, the absolute abundance of *P. copri* in feces collected from mice that had received *B. infantis* Bg2D9 was 3-orders of magnitude higher than in animals never exposed to *B. infantis* (**Extended Data Fig. 1c; Supplementary Table 5**).There was no statistically significant difference in weight gain from P23 to P42 between the mono- and bi-colonization groups, though interpreting this finding is at least partially confounded by the increased cecal size and fluid content observed in animals mono-colonized with *P. copri*.

The effects of *B. infantis* on *P. copri* did not generalize to *P. stercorea.* Unlike *P. copri*, the absolute abundance of *P. stercorea* in feces sampled on P21 and P24 was not significantly different in mice belonging to Arms 1 and 2 (*P*>0.05, Mann-Whitney U test). Prior to weaning at P24, the absolute abundance of *P. stercorea* was 5 orders of magnitude lower than that of *P. copri*. Throughout the post-weaning period, the absolute abundance of *P. stercorea* remained similar in members of both treatment arms (*P*>0.05, Mann-Whitney U test) but 2 orders of magnitude below that of *P. copri* (**Supplementary Table 3c**).

Principal components analysis (PCA) on the absolute abundance profiles of the 16 other organisms that had been administered to all three groups of mice revealed that over the course of the experiment *B. infantis* Bg2D9 colonization resulted in a fecal community composition that was distinct from that of animals colonized with *B. infantis* Bg463, regardless of their *Prevotella* colonization (**Fig. 1g**). In animals colonized with *Prevotella* and either of the two *B. infantis* strains (Arm 1 vs. Arm 2), these differences were observed as early as P21 (*P*=0.008; PERMANOVA) and became more pronounced by the end of the experiment at P53 (*P*<0.001; PERMANOVA) (**Fig. 1g**). Animals harboring *Prevotella*-containing communities, colonization with *B. infantis* Bg2D9 compared to Bg463 significantly increased the fitness of three organisms in the P53 fecal community and five organisms in the P53 cecal community (*P*<0.05, Kruskal-Wallis and post-hoc Dunn’s test with Bonferroni correction) while the absolute abundances of the other community members were not significantly affected (**Extended Data Fig. 2a,b; Supplementary Table 3c,d**). Surprisingly, only one significant difference, involving *B. catenulatum*, was apparent in fecal communities at P21 when *B. infantis* Bg2D9 achieved its highest abundance, suggesting that other factors continue to affect each organism’s fitness during the weaning transition and through to the end of the experiment (**Extended Data Fig. 2c; Supplementary Table 3c**). In contrast, in animals colonized with *B. infantis* Bg463, addition of *Prevotella* to the community did not result in significant differences in community composition at either P21 or P53, and only significantly increased the fitness of one organism (*S. gallolyticus*; see **Fig. 1g**, **Extended Data Fig. 2a**). However, when comparing Arms 1 and 3, the combination of *B. infantis* Bg2D9 and *Prevotella* colonization increased the fitness of a larger set of seven and six organisms in P53 fecal and cecal communities, respectively, including *B. catenulatum*, *B. obeum*, and *M. multacida -* three of the four organisms predicted to be capable of utilizing arabinose (**Extended Data Fig. 2a,b**); this suggests a potential synergistic interaction between the two organisms in mediating effects on community structure. Based on these results, we concluded that in the context of this preclinical model, (i) *B. infantis* Bg2D9 colonization was an important determinant of microbial community structure, including the fitness of *P. copri*; (ii) communities containing *B. infantis* 2D9 were associated with augmented weight gain, and (iii) the temporal profile of community member fitness produced when *B. infantis* 2D9 was included more closely resembled that of children in the clinical study who, during the weaning period when MDCF-2 treatment was initiated, all had substantial levels of *P. copri*^1^.

#### Metabolism of MDCF-2 glycans

Given these observed differences in microbial community structure, we used ultra-high performance liquid chromatography-triple quadrupole mass spectrometric (UHPLC-QqQ-MS)-based measurements of monosaccharide and linkage content of glycans to analyze metabolism of MDCF-2 and to determine if there were differences in carbohydrate utilization strategies between the groups. In our companion study^1^, we report increased levels of enzyme-resistant arabinose linkages, such as 5-Ara*f*, 2-Ara*f*, and 2,3-Ara*f*, in the feces of MDCF-2 treated children in the upper-compared to lower-quartile of WLZ responses. The results suggested increased consumption and preferential/partial degradation of MDCF-2 and arabinose-containing complementary foods among these MDCF-2 responders. Compared to the human study, our reverse translation experiments conducted in gnotobiotic mice provided a more controlled and sensitive environment for assaying the effects of *P. copri* on MDCF-2 glycan metabolism. We had access to and focused on the mouse cecum because we wanted to compare microbial gene expression with polysaccharide degradative capacity in a large gut habitat specialized for microbial fermentation^17^. Moreover, a uniform diet, comprised only of MDCF-2, could be monotonously administered to mice, eliminating potential confounding effects of variations in background diet encountered in humans.

While *B. infantis* Bg2D9 colonization seemed to be an important determinant of microbial community structure, *Prevotella* colonization drove degradation of MDCF-2 glycans, regardless of *B. infantis* colonization. *Prevotella* colonization significantly reduced the levels of arabinose in cecal glycans (*P*<0.01, Kruskal-Wallis and post-hoc Dunn’s test with Bonferroni correction; **Fig. 2a**) and levels of the arabinose-containing linkages t-Ara*p*, t-Ara*f*, 2-Ara*f*, 2,3-Ara*f*, and 3,4-Xyl*p*/3,5-Ara*f* (*P*<0.05, Kruskal-Wallis and post-hoc Dunn’s test with Bonferroni correction; **Fig. 2b**). These differences are supported by the fact that *P. copri* Bg131, unlike *P. stercorea*, contains PULs involved in processing arabinose-containing MDCF-2 glycans: *i.e.*, PUL27b specifies CAZymes known or predicted to digest arabinogalactan, while PUL2 possesses a fucosidase that could target the terminal residues found in arabinogalactan II (**Supplementary Table 1f**). In contrast, there were no significant differences between animals colonized with *B. infantis* Bg2D9 versus *B. infantis* Bg463 in Arms 1 and 2 for any of the monosaccharides or linkages measured (**Supplementary Table 6**). This is consistent with the low abundance of both *B. infantis* strains by the end of the experiment (**Fig. 1e**), suggesting that its influence on the fitness and carbohydrate utilization of other organisms is exerted earlier in the course of community assembly. Together, these results suggest that *Prevotella*-containing communities exhibit more complete degradation of branched arabinans and a greater degree of liberation of arabinose from MDCF-2 glycans.

**Fig. 2.**
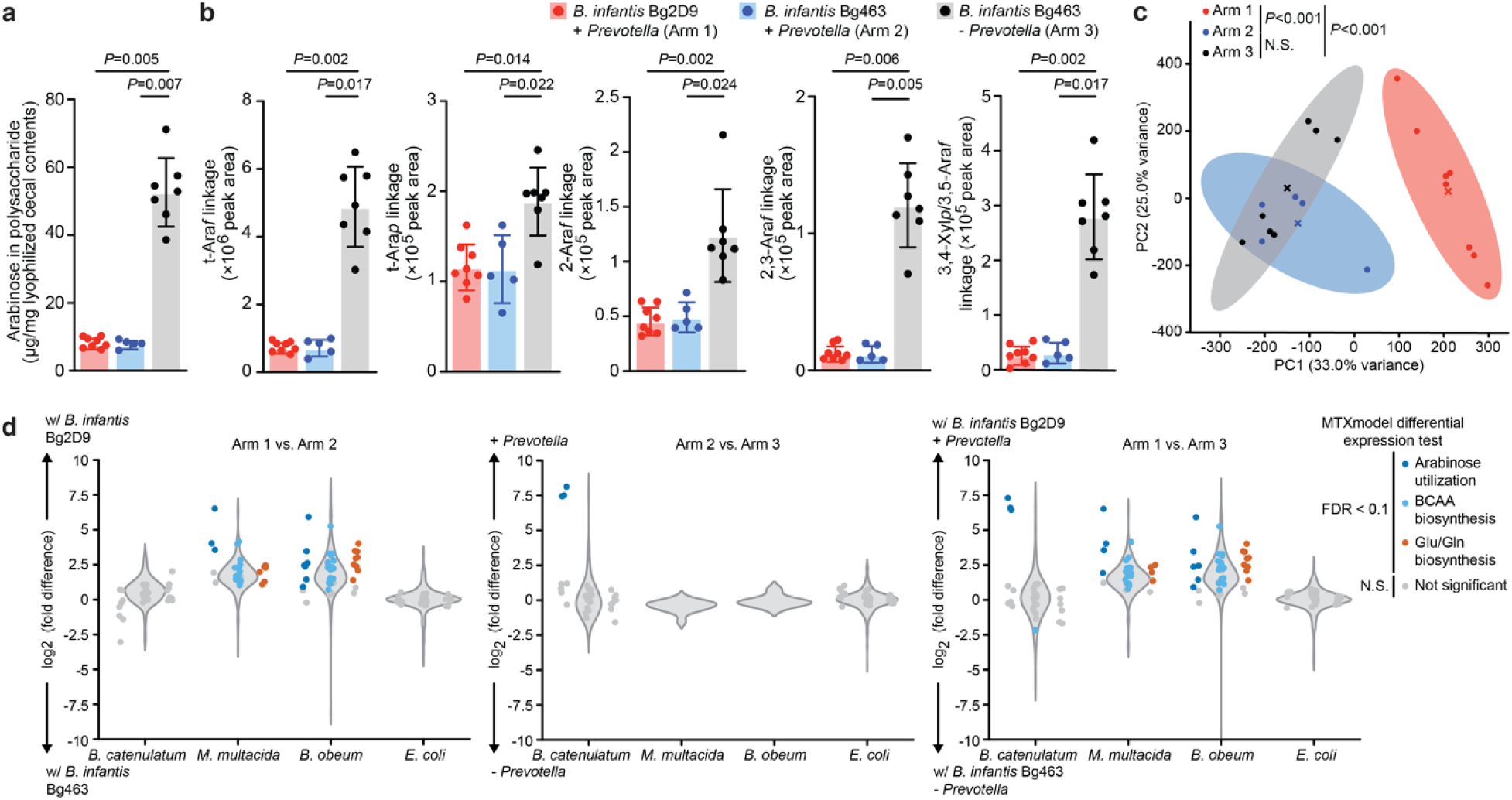
Targeted mass spectrometric and microbial RNA-Seq analyses of consortia of cultured age-discriminatory and WLZ-associated bacteria strains that colonized gnotobiotic mice. Cecal contents collected at the end of the experiment described in Fig. 1a were analyzed. **(a,b)** UHPLC-QqQ-MS-based quantitation of levels of total arabinose (panel a) and arabinose-containing glycosidic linkages (panel b) in cecal glycans collected at P53. Abbreviations: Ara*f*, arabinofuranose; Ara*p,* arabinopyranose; Xyl*p*, xylopyranose. Each dot represents an individual animal (n=5-8 mice/treatment arm). Mean values ± SD are shown. *P*-values were calculated by the Kruskal-Wallis test followed by post-hoc Dunn’s test with Bonferroni correction for panels a and b. (**c**) PCA of profiles of normalized meta-transcriptomic counts (see *Methods*). Centroids are denoted by a colored “X” for each group. *P*-values were calculated by PERMANOVA. (**d**) *MTXmodel* abundance-normalized differential expression analysis of genes involved in specific carbohydrate utilization and amino acid biosynthetic pathways in the four arabinose-utilizing bacteria. Violin plots show the distribution of log_2_ fold-differences for all expressed genes with metabolic pathway annotations in the indicated organism. Dots in panel d represent differential expression test results for individual genes involved in the corresponding pathway and are colored if their Benjamini-Hochberg adjusted *P*-value is less than 0.1. Abbreviations: BCAA, branched-chain amino acid; Glu, glutamate; Gln, glutamine.

#### Microbial community gene expression

To investigate how these changes in glycan utilization are associated with the expressed metabolic functions of community members, we performed microbial RNA-seq on cecal contents collected at the time of euthanasia (**Supplementary Table 7**). We first analyzed the expression levels of PULs by *P. copri* and *P. stercorea* to determine their individual contributions to the observed glycan utilization patterns. In both *Prevotella*-containing arms (Arms 1 and 2), *P. copri* PULs with predicted targets of starch and arabinogalactan (PUL 27a and 27b, respectively) were the most significantly enriched for higher levels of expression by GSEA [see *Methods*] (**Extended Data Fig. 3a; Supplementary Table 7b**).

In contrast to *P. copri*, only two *P. stercorea* PULs with predicted targets of α-mannan and *N*-linked glycans (PULs 5 and 7, respectively) were significantly enriched for higher expression (**Extended Data Fig. 3b; Supplementary Table 7b**). These two PULs had lower levels of expression than the *P. copri* PULs, consistent with the lower abundance of *P. stercorea* in the community. Furthermore, unlike the observed reductions in the arabinose content of cecal contents harvested from mice colonized with the *Prevotella-*containing consortia, there were no significant differences in mannose, N-acetylglucosamine, or N-acetylgalactosamine, the primary components of glycans targeted by these *P. stercorea* PULs (*P*>0.05; Kruskal-Wallis test; **Extended Data Fig. 3c**). These results indicate that *P. copri,* not *P. stercorea*, is responsible for increased liberation of arabinan from MDCF-2 and contributes to degradation of polysaccharides represented in MDCF-2.

We then turned to rest of the community to determine whether these changes in glycan metabolism had effects on the metabolic responses of other organisms. To assess the composition of expressed functions relative to the abundance of these organisms in the community, we performed PCA on meta-transcriptomic counts (reads from microbial RNA-Seq) normalized by paired metagenomic counts (reads from shotgun sequencing of cecal DNA) for genes originating from the 16 organisms that colonized all mice in all three arms of the experiment (i.e., the organisms that were not *B. infantis* or *Prevotella*). *B. infantis* Bg2D9 produced a meta-transcriptome that was distinct from both of the *B. infantis* Bg463-containing meta-transcriptomes (*P*<0.001, PERMANOVA; **Fig. 2c**). To identify metabolic responses of individual organisms to the presence or absence of either *B. infantis* Bg2D9 or *P. copri*, we performed differential expression testing incorporating normalization for underlying DNA abundances using *MTXmodel*^18^ to minimize the confounding effects of differential abundance on meta-transcriptomic counts. Differentially expressed genes associated with mcSEED metabolic pathway reconstructions are provided in **Supplementary Table 7c** and summarized in **Fig. 2d**. GSEA results for metabolic pathways enriched for increased or decreased expression in each comparison are provided in **Supplementary Table 7d**. Consistent with the UHPLC-QqQ-MS-based analysis of cecal glycans, arabinose utilization was among the most upregulated pathways in *B. catenulatum*, *M. multacida*, and *B. obeum* - three of the four organisms predicted to be capable of utilizing arabinose (**Fig. 2d; Supplementary Table 7d**). These three organisms were also significantly more abundant with *B. infantis* Bg2D9 colonization (**Extended Data Fig. 2b**). *B. catenulatum* upregulated arabinose utilization genes directly in response to *P. copri* colonization (Arm 2 vs Arm 3; **Fig. 2d**). In contrast, *M. multacida* and *B. obeum* demonstrated upregulation of arabinose utilization in response to *B. infantis* colonization (Arm 1 vs Arm 2; **Fig. 2d**), though these changes are likely driven in part by their dependence on *B. infantis* Bg2D9 for colonization. *M. multacida* and *B. obeum* also demonstrated significant upregulation of almost all their genes involved in biosynthesis of the branched chain amino acids as well as glutamate and glutamine with *B. infantis* Bg2D9 colonization (**Fig. 2d; Supplementary Table 7d**). Together, these results suggest that while *B. infantis* Bg2D9 colonization affects the abundances of these organisms in the community, glycosidic activities (e.g., arabinan degradation) associated with *P. copri* colonization are a primary determinant of their metabolic responses.

### snRNA-Seq of intestinal gene expression

We next tested whether the respective effects of *B. infantis* Bg2D9 and *P. copri* on microbial community structure and expressed metabolic functions were associated with metabolic changes in epithelial cells in portions of the small intestine that are dedicated to nutrient absorption. Given that (i) the combination of *B. infantis* Bg2D9 and *P. copri* mediated a set of effects greater than those mediated by either organism alone and (ii) successful colonization with both *B. infantis* and *P. copri* before and through the weaning transition at P21 in Arm 1 better represented the microbial communities of children in the clinical study^1^, we advanced small intestinal samples from Arms 1 and 3 for further analysis. Because of the different *B. infantis* strains used and the greater fitness and greater expression of PULs targeting MDCF-2 glycans exhibited by *P. copri* but not *P. stercorea*, we refer to Arm 1 as ‘with *B. infantis* Bg2D9 and with *P. copri’* and Arm 3 as ‘with *B. infantis* Bg463 and without *P. copri*’ for these comparisons of the effect of the combination of these two organisms.

Histomorphometric analysis of villus height and crypt depth in jejunums harvested from mice in the ‘with *B. infantis* Bg2D9 and with *P. copri’* and ‘with *B. infantis* Bg463 and without *P. copri*’ groups (n=8 and 7, respectively) disclosed no statistically significant architectural differences between the two treatment groups (*P*>0.05; Mann-Whitney U test; **Supplementary Table 8**). We subsequently turned to snRNA-Seq to investigate whether these two colonization states produced differences in expressed functions along the crypt-villus axis in jejunal tissue collected from P53 animals (n=4/treatment arm; **Fig 3a-c; Extended Data Fig. 4; Supplementary Table 9**). A total of 30,717 nuclei passed our quality metrics (see *Methods*). Marker gene-based annotation disclosed cell clusters that were assigned to the four principal intestinal epithelial cell lineages (enterocytic, goblet, enteroendocrine, and Paneth cell) as well as to vascular endothelial cells, lymphatic endothelial cells, smooth muscle cells and enteric neurons (**Extended Data Fig. 4a,b**). Marker gene analysis allowed us to further subdivide the enterocytic lineage into three clusters: ‘villus-base’, ‘mid-villus’ and ‘villus-tip’. Pseudobulk snRNA-seq analysis, which aggregates transcripts for each cell cluster and then uses edgeR to identify differentially expressed genes in each cluster^19,20^, disclosed that a majority of all statistically significant differentially expressed genes (3,651 of 5,765; 63.3%) were assigned to the three enterocyte clusters (**Extended Data Fig. 4c; Supplementary Table 9b**).

**Fig. 3.**
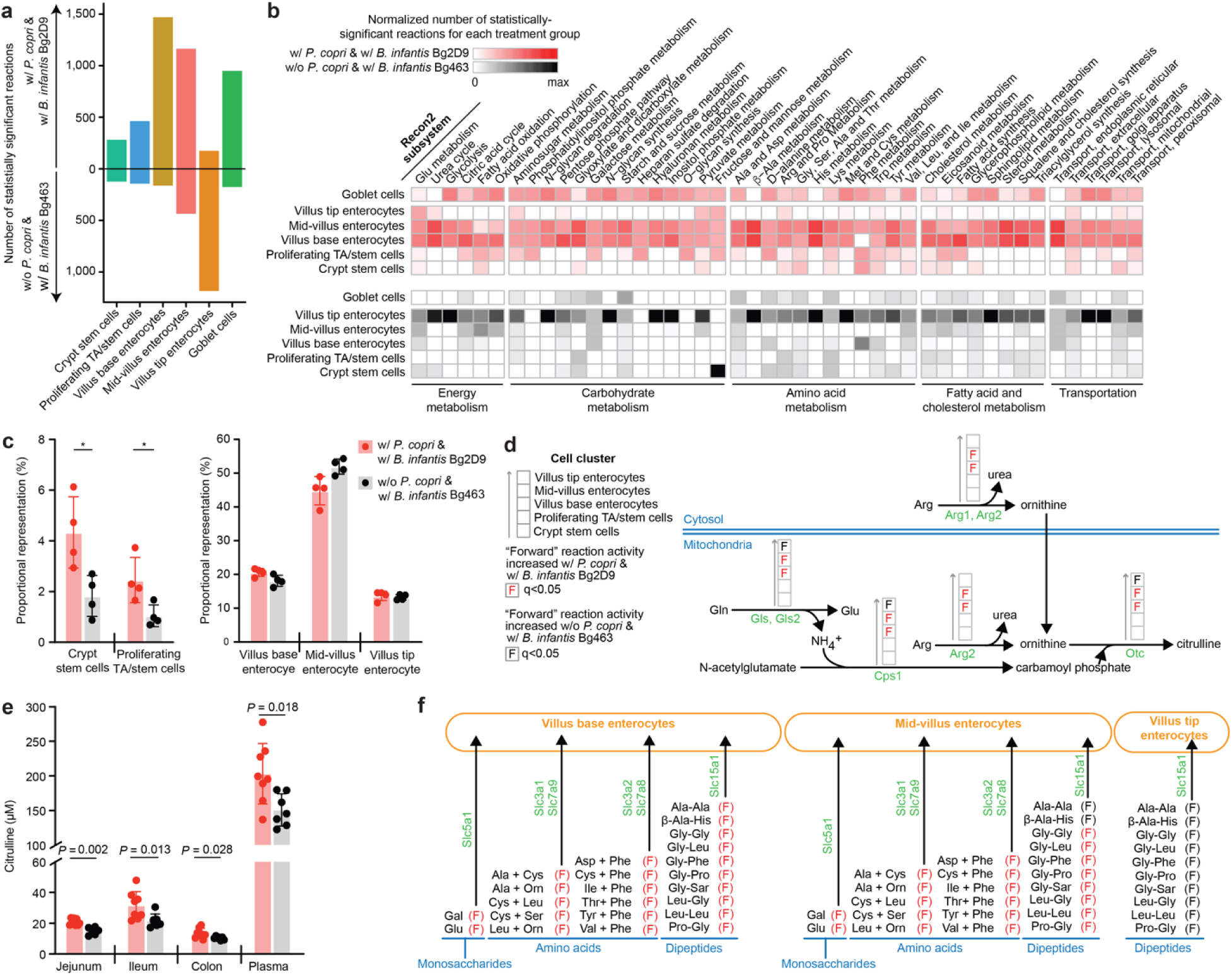
snRNA-Seq analysis and targeted mass spectrometric analysis of intestinal tissue and plasma harvested from mice containing bacterial communities with or without *P. copri* and two different strains of *B. infantis*. Jejunal tissue samples collected from Arm 1 (w/ *P. copri* & w/ *B. infantis* Bg2D9) and Arm 3 (w/o *P. copri* & w/ *B. infantis* Bg463) at the end of the experiment (P53) described in Fig. 1a were analyzed (n=4 samples/treatment arm). (**a**) The number of Recon2 reactions with statistically significant differences in their predicted flux between mice in Arm 1 and Arm 3. (**b**) The number of Recon2 reactions in each Recon2 subsystem that are predicted to have statistically significant differences in their activities between the two treatment groups. Colors denote values normalized to the sum of all statistically significantly different Recon2 reactions found in all selected cell clusters for a given Recon2 subsystem in each treatment group. (**c**) Proportional representation of cell clusters identified by snRNA-Seq. Asterisks denote ‘statistically credible differences’ as defined by *scCODA* (see **Supplementary Table 9c** and *Methods*). (**d**) Selected Recon2 reactions in enterocyte clusters distributed along the villus involved in the urea cycle and glutamine metabolism. (**e**) Targeted mass spectrometric quantifications of citrulline levels along the length of the gut and in plasma. Mean values ± SD and *P*-values from the Mann-Whitney U test are shown. Each dot represents an individual animal (n=7-8). (**f**) Effect of colonization with bacterial consortia containing or lacking *P. copri* on extracellular transporters for monosaccharides, amino acids and dipeptides. Sar: sarcosine. These transporters were selected and the spatial information of their expressed region along the length of the villus was assigned based on published experimental evidence^34^. Arrows in panels b and e indicate the “forward” direction of each Recon2 reaction. The Wilcoxon Rank Sum test was used to evaluate the statistical significance of the net reaction scores (panel a, b, d and e) between the two treatment groups. *P*-values were calculated from Wilcoxon Rank Sum tests and adjusted for multiple comparisons (Benjamini-Hochberg method); a q-value < 0.05 was used as the cut-off for statistical significance.

We initially used *NicheNet*^21^ to evaluate the effects of the *B. infantis* Bg2D9- and *P. copri* Bg131-containing community on intercellular communications. *NicheNet* integrates information on signaling and gene regulation from publicly available databases to build a “prior model of ligand-target regulatory potential” and then predicts potential communications between user-defined “sender” and “receiver” cell clusters ^21^. After incorporating snRNA-Seq-based expression data from both sender and receiver cells, *NicheNet* computes a list of potential ligand-receptor interactions between senders and receivers. The ligand-receptor interactions in the resulting list are then ranked based on their effect on expression of downstream genes in their signaling pathway (*i.e.*, more downstream genes are expressed in a ‘high-ranking’ interaction). After this ranking step, an additional filter is applied, with ligand-receptor interactions having firm experimental validation in the literature designated as “bona fide” interactions. Finally, *NicheNet* uses information generated by Seurat^22^ from a snRNA-Seq dataset to identify altered “bona fide” ligand-receptor interactions.

We designated six of the epithelial cell clusters (crypt stem cells, proliferating transit amplifying (TA)/stem cells, villus base, mid-villus, and villus tip enterocytes and goblet cells) as “receiver cells” while all clusters (both epithelial and mesenchymal) were designated “sender cells”. We then conducted *NicheNet* analysis for each sender-receiver pair. **Extended Data Fig. 5** shows bona fide ligand-receptor interactions that are altered between the two colonization conditions for each receiver cell cluster.

Ligands identified include those known to affect cell proliferation (*igf-1*), cell adhesion (*cadm1, cadm3*, *cdh3*, *lama2*, *npnt*), zonation of epithelial cell function/differentiation along the length of the villus (*bmp4*, *bmp5*), as well as immune responses (*cadm1*, *il15*, *tgfb1*, *tnc*) (**Extended Data Fig. 5**). Among all receiver cell clusters, crypt stem cells exhibited the highest number of altered bona fide ligand-receptor interactions. For example, Igf-1 is known to enhance intestinal epithelial regeneration^23^. We found that colonization with the *P. copri-*containing consortium was associated with markedly elevated expression of *igf-1* in goblet and lymphatic endothelial sender cells that signals to crypt stem cell receivers.

We subsequently applied the *Compass* algorithm^24^ to our snRNA-Seq datasets to generate *in silico* predictions of the effects of the consortia containing the combination of *B. infantis* Bg2D9 and *P. copri* on the metabolic states of (i) stem cell and proliferating TA cell clusters positioned in crypts of Lieberkühn, (ii) the three villus-associated enterocyte clusters, and (iii) the goblet cell cluster. *Compass* combines snRNA-seq data with the Recon2 database ^25^. This database describes 7,440 metabolic reactions grouped into 99 Recon2 subsystems^25^, plus information about reaction stoichiometry, reaction reversibility, and associated enzyme(s). Using snRNA-seq data, *Compass* computes a score for each metabolic reaction. If the metabolic reaction is reversible, then one score is calculated for the “forward” reaction and another score is calculated for the “reverse” reaction^25^. We calculated a ‘metabolic flux difference’ (see *Methods*) to quantify the difference in net flux for a given reaction (*i.e.*, the forward and reverse activities) between the two treatment groups.

**Fig. 3** shows the predicted metabolic flux differences for Recon2 reactions in enterocytes distributed along the length of the villus and in goblet cells. In clusters belonging to the enterocyte lineage, the number of statistically significant differences is greatest in villus base enterocytes and decreases towards the villus tip (**Fig. 3a**). Mice in the ‘w/ *P. copri* & w/ *B. infantis* Bg2D9’ treatment group, relative to their ‘w/o *P. copri* & w/ *B. infantis* Bg463’ counterparts, had the greatest predicted increases in the activities of subsystems related to energy metabolism, the metabolism of carbohydrates, amino acids and fatty acids, as well as various transporters, in villus base and mid-villus enterocytes (**Fig. 3b**, **Extended Data Fig. 6**).

While enterocytes prioritize glutamine as their primary energy source, they are also able to utilize fatty acids and glucose. The *Compass*-defined increase in reactions related to fatty acid oxidation that occur in the villus enterocytes of mice in the Arm 1 ‘w/ *P. copri* & w/ *B. infantis* Bg2D9’ group compared to those in the Arm 3 ‘w/o *P. copri* & w/ *B. infantis* Bg463’ group extended to their crypts of Lieberkühn (**Fig. 3b**). Fatty acid oxidation has been linked to intestinal stem cell maintenance and regeneration^26^. Mice colonized with *P. copri* and *B. infantis* Bg2D9 exhibited ‘statistically credible increases’ in the proportional representation of crypt stem cells and proliferating TA/stem cells but not in their villus-associated enterocytic clusters (**Fig. 3c**; also see **Supplementary Table 9c** for results regarding all identified epithelial and mesenchymal cell clusters). [The term ‘statistically credible difference’ was defined by *scCODA*^27^ (see *Methods*)]. Compared to mice colonized with *B. infantis* Bg463 and lacking *P. copri*, those colonized with *B. infantis* Bg2D9 and *P. copri* also had predicted increases in energy metabolism in their goblet cells, as judged by the activities of subsystems involved in glutamate (Glu) metabolism, the urea cycle, fatty acid oxidation and glycolysis (**Fig. 3b**).

Citrulline is generally poorly represented in human diets; it is predominantly synthesized via metabolism of glutamine in small intestinal enterocytes and transported into the circulation^28^. Both glutamate and arginine are important for citrulline production in enterocytes^29^. Glutaminase (Gls) and glutamate dehydrogenase (GluD) in the glutamine pathway provide ammonia for generating carbamoyl phosphate (**Fig. 3d**), while arginine is a primary precursor for ornithine synthesis^30^. Ornithine transcarbamylase (Ots) produces citrulline from carbamoyl phosphate and ornithine. *Compass* predicted that mice harboring *B. infantis* Bg2D9 and *P. copri* exhibit statistically significant increases in these reactions in their villus base and mid-villus enterocyte clusters [q<0.05 (adjusted *P-*value); Wilcoxon Ranked Sum test; **Fig. 3d**]. Follow-up targeted mass spectrometric analysis confirmed that citrulline was significantly increased in jejunal, ileal and colonic tissue segments, as well as in the plasma of mice in Arm 1 compared to Arm 3 (*P*<0.05; Mann-Whitney U test; **Fig. 3e**). Notably, studies of various enteropathies and short bowel syndrome have demonstrated that citrulline is a quantitative biomarker of metabolically active enterocyte mass and its levels in plasma are indicative of the absorptive capacity of the small intestine^28^. Citrulline has also been reported to be markedly lower in blood from children with severe acute malnutrition compared to healthy children^31^. Low plasma citrulline levels have also been described in cohorts of undernourished children with environmental enteric dysfunction and higher levels were predictive of future weight gain^32,33^.

The presence of *P. copri* and *B. infantis* Bg2D9 was also associated with significantly greater predicted activities in Recon2 subsystems involved in transport of nine amino acids (including the essential amino acids leucine, isoleucine, valine, and phenylalanine), dipeptides and monosaccharides (glucose and galactose) in villus base and mid-villus enterocytes (**Fig. 3f**). These predictions suggest a greater absorptive capacity for these important growth-promoting nutrients, which are known to be transported within the jejunum at the base and middle regions of villi^34^.

### Further assessment of the effects of *P. copri* on host metabolism and weight gain

Based on this first experiment, we concluded that *B. infantis* Bg2D9 could promote weight gain and affect microbial community structure, including *P. copri* colonization. Given that *P. copri* colonization was in turn linked to changes in glycan metabolism and microbial community gene expression, we next tested whether *P. copri* could play a role in mediating effects on host weight gain and intestinal metabolism that were observed with the combination of *B. infantis* Bg2D9 and *P. copri*. Therefore, we repeated the experiment described above with a larger number of animals (4 dually housed germ-free dams yielding 18-19 viable pups per arm). To examine whether the weight gain phenotype and metabolic alterations observed in the experiment described above could be attributed to the presence or absence of *P. copri* in the microbial community, we administered *B. infantis* Bg2D9 in both arms of this repeat experiment. Outside of this change, the same cultured strains, the same sequence of their introduction and the same sequence of diet switches were applied (**Extended Data Fig. 7a**). Reproducible colonization of consortium members within each arm was confirmed by quantifying their absolute abundances in cecal samples collected at the time of euthanasia (P53; see **Supplementary Table 10**). As in the previous experiment, animals in the arm containing *P. copri* exhibited significantly greater weight gain between P23 and P53 than those in the no *P. copri* arm [*P*<0.05; linear mixed-effects model (see *Methods*)] (**Extended Data Fig. 7b**).

#### Mass spectrometric analysis of host metabolism

We used targeted mass spectrometry to quantify levels of 20 amino acids, 19 biogenic amines, and 66 acylcarnitines in the jejunum, colon, gastrocnemius, quadriceps, heart muscle, and liver of the two groups of mice. Additionally, we quantified the 66 acylcarnitines in their plasma. The results are described in **Supplementary Table 11** and **Extended Data Fig. 7c-e**. Consistent with the previous experiment, citrulline, the biomarker for metabolically active enterocyte biomass, was significantly elevated in the jejunums of mice belonging to the w/ *P. copri* group (*P*<0.05; Mann-Whitney U test) (**Extended Data Fig. 7c**; **Supplementary Table 11a**).

We observed significant elevations of acylcarnitines derived from palmitic acid (C16:0), stearic acid (C18:0), oleic acid (C18:1), linoleic acid (C18:2), and linolenic acid (C18:3) in the jejunums of *P. copri*-colonized animals (*P*<0.01; Mann-Whitney U test) (**Extended Data Fig. 7d**); these are the major fatty acids found in soybean oil^35^, which is the principal source of lipids in MDCF-2. These acylcarnitine chain lengths were found at higher abundance than all other medium or long-chain acylcarnitine species in our jejunal samples, indicating their role as primary dietary lipid energy sources (**Supplementary Table 11b)**. Elevation of these species suggests increased transport and ß-oxidation of long-chain dietary lipids in the jejunums of the *P. copri-*colonized animals.

Analysis of colonic tissue showed significant elevation of C16:0, C18:1, and C18:2 acylcarnitines in *P. copri*-colonized animals, suggesting that β-oxidation is also elevated in tissue compartments not directly involved in lipid absorption (*P*<0.01; Mann-Whitney U test) (**Extended Data Fig. 7e**; **Supplementary Table 11b**). This finding was matched by a significant elevation in plasma levels of non-esterified fatty acids in w/ *P. copri* animals, suggesting higher circulation of dietary lipids which would support fatty acid β-oxidation in peripheral tissues (*P*<0.05; Mann-Whitney U test) (**Extended Data Fig. 7f**; **Supplementary Table 11c**). We extended our targeted LC-MS to liver, gastrocnemius muscle, quadriceps and heart. The only statistically significant difference in levels of acylcarnitines whose chain length corresponded to components of soybean oil was an increase in 18:2 and 18:3 species in the myocardium of mice with *P. copri* compared to those lacking this organism (**Supplementary Table 11b**). Additionally, jejunal levels of C3 and C4 acylcarnitines as well as colonic levels of C4 and C5 acylcarnitines known to be derived from branched-chain amino acid catabolism, were significantly elevated in the *P. copri*-colonized animals (*P*<0.05; Mann-Whitney U test; **Extended Data Fig. 7d,e** and **Supplementary Table 11b**).

### *P. copri* colonization and weight gain in another diet context

The focus of these ‘reverse translation’ experiments in gnotobiotic mice was to establish whether *P. copri* MAGs identified in the human study play a key role in the metabolism of MDCF-2 glycans and host responses. As such, our mouse experiments examined the effects of *P. copri* Bg131 in the context of the MDCF-2 diet. The experiment shown in **Extended Data Fig. 8** illustrates the challenge posed in testing the effects of *P. copri* colonization in the context of a ‘control’ diet representative of that typically consumed by 18-month-old Bangladeshi children living in Mirpur (‘Mirpur-18 diet’)^5^. The design was similar to that employed for the experiments described in **Fig. 1** and **Extended Data Fig. 7** with two exceptions; (i) *B. infantis* Bg2D9 was used in both groups and (ii) on P24 pups from different litters were mixed and randomly assigned to two diet treatment groups, MDCF-2 and Mirpur-18 (n=2 dams and 12 pups per group). The absolute abundances of community members were quantified in cecal contents harvested at the time of euthanasia on P53. While the absolute abundance of *P. copri* Bg131 was not significantly different between the two diet groups (*P*>0.05; Mann-Whitney), there were statistically significant differences between the abundances of 11 of the 19 community members (**Extended Data Fig. 8b,c**). Nonetheless, we proceeded to test whether the increased weight gain phenotype associated with the presence of *P. copri* in the community was evident in the Mirpur-18 diet context. To do so, we repeated the dam-to-pup microbial transmission experiment where all animals were weaned onto the Mirpur-18 diet but where one group had received *P. copri* Bg131 (n=8 animals) and the other had not (n=9). All animals were euthanized on P53. *P. copri* successfully colonized mice and was maintained throughout the experiment at levels comparable to previous experiments (10.4±0.1 log_10_ genome equivalents per gram cecal contents at P53). Importantly, there was no statistically significant difference in weight gain between the two groups [*P*=0.297; linear mixed-effects model (see *Methods*)]. These findings provide evidence that the effect of *P. copri* on weight gain in this preclinical model is diet dependent.

### Functional evaluation of cultured *P. copri* strains more closely resembling MAGs Bg0018 and Bg0019

To further test the causal relationship between *P. copri* MAGs Bg0018/Bg0019, glycan and host metabolism, and weight gain in the context of MDCF-2, we turned to cultured *P. copri* strains whose genomic features were closer to the MAGs than those represented in strain Bg131. *P. copri* Bg131 shared notable but incomplete conservation of PULs and other genomic features represented in MAGs Bg0018 and Bg0019. We have previously characterized five additional fecal *P. copri* strains that we cultured from Bangladeshi children living in Mirpur^1^. Two of these strains (BgD5_2 and BgF_2), had greater genomic similarity to MAGs Bg0018 and Bg0019 than the other isolates, including *P. copri* Bg131, as quantified by phylogenetic distance and PUL content. Nine of the 10 functionally conserved PULs in Bg0018/Bg0019 were present in *P. copri* BgD5_2 and BgF5_2 compared to five in the case of *P. copri* Bg131 (**Supplementary Table 12a,b**; **Fig. 4a**). All 103 genes in these nine conserved PUL homologs shared 100% nucleotide sequence identity between the two isolates. Moreover, of the 198 genes comprising the 22 PULs contained in each isolate’s genome, 195 shared 100% nucleotide sequence identity (**Supplementary Table 12c**). *In silico* metabolic reconstructions disclosed that both the BgD5_2 and BgF5_2 strains shared 103/106 (97%) binary phenotypes with MAG Bg0018 and 102/106 (96%) with MAG Bg0019, including 53/55 (96%) and 54/55 (98%) of their carbohydrate utilization pathways with MAGs Bg0018 and Bg0019, respectively (**Supplementary Table 12d**). In addition, results of *in vitro* growth assays conducted in defined medium supplemented with different glycans represented in MDCF-2 disclosed that strain BgF5_2 exhibited stronger preference than Bg131 for glycans enriched in and/or unique to MDCF-2 compared to RUSF (i.e., arabinan, arabinoxylan, galactan, galactomannan)^1^.

**Fig. 4.**
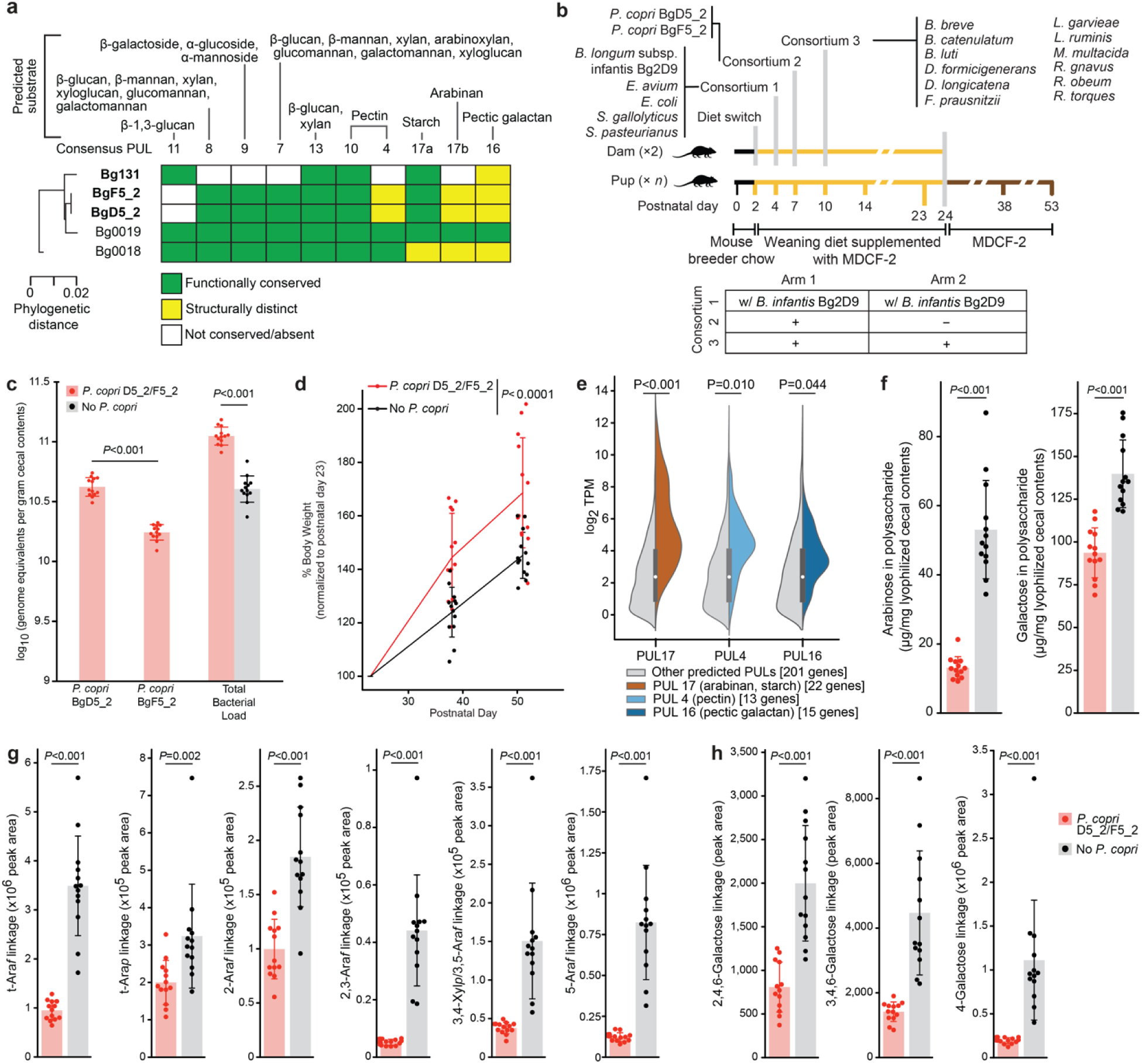
Testing the effects of pre-weaning colonization with two *P. copri* strains closely related to MAGs Bg0018 and Bg0019 on host weight gain and MDCF-2 glycan degradation. (**a**) Comparison of polysaccharide utilization loci (PULs) highly conserved in the two *P. copri* MAGs with their representation in the three cultured *P. copri* strains. (**b**) Study design (n=2 dams and 13 offspring/ treatment arm). (**c**) Absolute abundance of *P. copri* strains and total bacterial load in cecal contents collected at the end of the experiment (P53). (**d**) Body weights of the offspring of dams, normalized to postnatal day 23 [linear mixed effects model (see *Methods*)]. (**e**) GSEA of expression of PULs shared by *P. copri* BgD5_2 and BgF5_2 in the cecal contents of animals. Adjusted *P*-values were calculated using GSEA ranking genes by their mean log_2_ transcripts per million (TPM) across the *P. copri* colonized samples, with each PUL comprising a gene set against the background of all predicted PUL genes. Violin plots show the log_2_ TPM of all genes assigned to any of the 22 predicted PULs in each isolate (n=201 genes) in each of the samples, split to show homologs of consensus PUL 17 (arabinan, starch; n=22 genes), PUL 4 (pectin; n=13 genes), and PUL 16 (pectic galactan; n=15 genes) in color compared to the remainder of all PUL genes in gray. Internal box plots show the median (circle) and quartiles (thick line) for all genes assigned to PULs. (**f**) UHPLC-QqQ-MS analysis of total arabinose and galactose in glycans present in cecal contents collected at P53. (**g,h**) UHPLC-QqQ-MS of glycosidic linkages containing arabinose (panel g) and galactose (panel h) in cecal contents. Mean values ± SD are shown. *P-*values were calculated using a Mann-Whitney U test. Each dot in panels b-h represents an individual animal.

To directly determine whether pre-weaning colonization with *P. copri* strains resembling MAGs Bg0018 and Bg0019 is sufficient to promote growth and produce the metabolic effects described above, we performed an experiment whose design (**Fig. 4b**) was similar to our previous experiments (**Fig. 1b** and **Extended Data Fig. 7a**), but with several modifications. *First*, because of their greater genomic similarity to WLZ-associated MAGs Bg0018 and Bg0019, we used *P. copri* BgD5_2 and BgF5_2 in place of *P. copri* Bg131. *Second*, to control for differences in *B. infantis* strain used in the initial experiment (**Fig. 1**), both arms received *B. infantis* Bg2D9 in this ‘third’ experiment (as was the case in the second experiment described in **Extended Data Fig. 7**). *Third*, because *P. stercorea* had colonized at a lower abundance than *P. copri* and did not express CAZymes related to MDCF-2 glycans, it was not included in the second gavage mixture, which now only contained *P. copri*. *Fourth*, given that *B. infantis* Bg2D9 promoted pre-weaning colonization of *P. copri* in the initial experiment, we omitted the fourth gavage, previously administered at the end of the weaning period, that had included *P. copri* and *P. stercorea*. As before, the control group of animals were those that did not receive *P. copri* (n=2 dams and 13 pups/treatment group).

Shotgun sequencing of DNA isolated from cecal contents collected at the time of euthanasia (P53) confirmed that animals in the experimental group had been colonized with both *P. copri* isolates as well as all other members of the defined consortia (**Supplementary Table 13**). Even though strain BgD5_2 grew much more poorly than BgF5_2 when cultured in defined medium^1^, in animals colonized with both isolates, the BgD5_2 strain was present at higher absolute abundance than the BgF5_2 strain (**Fig. 4c**); their relative abundances were 37.8±4.4% and 15.5±1.0%, respectively, compared with 31±6.6% and 24±8.0% for *P. copri* Bg131 in the experiments described in **Fig. 1b** and **Extended Data Fig. 7a**. Comparing the two groups disclosed that colonization with BgD5_2 and BgF5_2 augmented community biomass without displacing other bacteria (**Fig. 4c; Supplementary Table 13b**).

We observed a significantly greater increase in body weight between P23 and P53 in mice colonized with *P. copri* BgD5_2 and BgF5_2 compared to those without *P. copri* [*P*<0.0001; linear mixed-effects model] (**Fig. 4d)**. The difference in the mean percent increase in postweaning weight between the experimental and control groups (24%) was comparable to that documented in the two previous experiments shown in **Fig. 1b** and **Extended Data Fig. 7a** (25% and 13% respectively). As in these previous experiments, the weight difference was not attributable to differences in cecal size.

Mass spectrometry confirmed that preweaning colonization with *P copri* affected intestinal lipid metabolism and was a major determinant of MDCF-2 glycan degradation. Targeted LC-MS of ileal and colonic tissue revealed a significant elevation of long-chain acylcarnitines corresponding to soybean oil lipids (**Extended Data Fig. 9; Supplementary Table 14**), consistent with changes observed in the experiment described in **Extended Data** Fig. 7. Microbial RNA-Seq of cecal contents revealed that among all PUL genes, those present in the three conserved PULs with predicted arabinan/starch, pectin, and pectic galactan substrates were significantly enriched for higher levels of expression (See *Methods*) (**Fig. 4e; Supplementary Table 15**). UHPLC-QqQ-MS of monosaccharides in glycans present in cecal contents indicated that the presence of *P. copri* BgD5_2 and F5_2 resulted in significantly lower levels of arabinose, consistent with our previous observations using *P. copri* Bg131, as well as galactose (a finding specific to this experiment) (**Fig. 4f; Supplementary Table 16**). Colonization with *P. copri* BgD5_2 and BgF5_2 also significantly lowered levels of all arabinose-containing glycosidic linkages measured, as well as three galactose-containing linkages (**Fig. 4g,h; Supplementary Table 16**). Together, these data indicate that the PUL content of these two isolates is associated with enhanced degradation of MDCF-2 glycans compared to the *P. copri* Bg131-containing microbial community. Targeted UPHLC-QqQ-MS measurements of all 20 amino acids and seven B vitamins also revealed that compared to the control group, colonization with these strains was associated with significantly higher cecal levels of two essential amino acids (tryptophan, lysine), seven non-essential amino acids (glutamate, glutamine, aspartate, asparagine, arginine, proline, glycine) and pantothenic acid (vitamin B5) (**Supplementary Table 17**).

Based on all of these experiments, we concluded that (i) pre-weaning colonization with *P. copri* augments weight gain in the context of the MDCF-2 diet, (ii) the presence of specific strains of this species is a major determinant/effector of MDCF-2 glycan degradation, and (iii) incorporating these strains into the gut community changes intestinal cellular metabolism.

## Discussion

Accessing tissue from different regions of the human intestine, as well as from extra-intestinal sites, represents a major challenge when trying to characterize the mechanisms by which microbiome-targeted nutritional interventions impact the operations of microbial community members, and how these changes alter human physiology at a molecular, cellular and systems level. In this study, we illustrate a ‘reverse translation’ strategy that can be used to address this challenge. Gnotobiotic mice were colonized with defined consortia of age- and WLZ-associated bacterial strains cultured from fecal samples collected from children living in a Bangladeshi community where the prevalence of malnutrition is high. *Prevotella copri* was represented by cultured isolates whose genomic features, including PULs and metabolic pathways involved in carbohydrate utilization, are highly similar to MAGs associated with improved weight gain in the clinical trial. Dam-to-pup transmission of these communities occurred in the context of a sequence of diets that re-enacted those consumed by children enrolled in a clinical study of a microbiome-directed complementary food (MCDF-2). Microbial RNA-Seq and targeted mass spectrometry of glycosidic linkages present in intestinal contents provided evidence that *P. copri* plays a key role in the metabolism of polysaccharides contained in MDCF-2. Consistent with this, *P. copri-* associated weight gain in the preclinical model was dependent on consumption of MDCF-2; this phenotype was not observed when a diet commonly consumed by Bangladeshi children was administered instead, despite comparable levels of *P. copri* colonization in the two diet contexts. Single nucleus RNA-Seq and targeted mass spectrometry of the intestine indicated that colonization with the consortium that contains the combination of *B. infantis* Bg2D9 and *P. copri* increases the uptake and metabolism of lipids (including those fatty acids that are most prominently represented in the soybean oil that comprises the principal lipid component of MDCF-2). Additional effects on uptake and metabolism of amino acids (including essential amino acids) and monosaccharides were predicted and in select cases validated by mass spectrometric assays. These effects on nutrient processing and energy metabolism involve proliferating epithelial progenitors in the crypts as well as their descendant lineages distributed along the villus. snRNA-Seq revealed discrete spatial features of these effects, with populations of enterocytes positioned at the base-, mid- and tip regions of villi manifesting distinct patterns of differential expression of a number of metabolic functions.

Inspired by the results of the clinical trial, the goal of our reverse translation experiments was to ascertain the impact of the presence or absence of *P. copri* in a model that emulated postnatal gut microbial community assembly and exposure to MDCF-2. The current study raises several questions that have both mechanistic and therapeutic implications. We were not able to successfully mono-colonize mice with our cultured Bangladeshi *P. copri* strains. Therefore, we could not directly test the effects of these strains *in vivo* on MDCF-2 glycan metabolism, weight gain and/or gut epithelial biology in the absence of other potential microbial interactions. Additional work, involving systematic manipulation of the composition of the bacterial consortia introduced into dams (and subsequently transmitted to their offspring) will be required to ascertain the extent to which *P. copri* has (i) direct effects on intestinal epithelial gene expression and host metabolism versus (ii) effects of other community members that are dependent upon its presence or absence. If *P. copri* has direct effects on the host, it remains to be determined whether the mediators of these effects are the direct products of MDCF-2 glycan metabolism, or the products of other metabolic pathways in *P. copri* whose activities are regulated by biotransformation of these glycans, or other MDCF-2 components. Future studies are also needed to disambiguate the extent to which the observed effects of *P. copri* on gut epithelial carbohydrate, lipid and amino acid metabolism contribute to weight gain. The spatial features of metabolic pathway expression documented by snRNA-seq must be characterized further. This effort will be technically challenging; e.g., it could require (i) documenting the distribution of *P. copri* and other community members along the crypt-villus axis, (ii) advancing methods for spatial transcriptomics^34,36^ so that they can be (simultaneously) applied to both epithelial cell lineages and microbial community members, and (iii) using *in situ* mass spectrometry to directly characterize the metabolic profiles of discrete gut cell populations.

The relationship between prominent initial colonization by *B. infantis* Bg2D9 and the capacity of *P. copri* to subsequently colonize also needs further investigation. As noted above, *B. infantis* Bg2D9 contains several genomic loci not represented in most other cultured *B. infantis* strains, which could enhance its ability to utilize a variety of dietary carbohydrates^12^. In principle, these loci could support increased fitness of *B. infantis* Bg2D9 in malnourished children whose consumption of breast milk is low. Bangladeshi infants and young children with severe acute malnutrition (SAM) have markedly lower or absent levels of *B. infantis* compared to their healthy counterparts^12^. The *B. infantis*-*P. copri* interaction documented in this preclinical study provides a rationale for testing the effects of first administering *B. infantis* Bg2D9 to individuals with SAM and subsequently MDCF-2 to restore age-appropriate microbiome configurations and promote healthy growth.

In summary, this and our companion study^1^ illustrate one approach for identifying members of a gut microbial community that function as principal metabolizers of dietary components as well as key effectors of host biological responses. The results can provide a starting point for developing microbiome-based diagnostics for stratification of populations of undernourished children who are candidates for treatment with a given MDCF, and for monitoring their treatment responses including in adaptive clinical trial designs. Another potential return on investment for this approach is a knowledge base needed for (i) creating ‘next generation’ MDCFs composed of (already) identified bioactive components, but from alternative food sources that may be more readily available, affordable and culturally acceptable for populations living in different geographic locales; (ii) making more informed decisions about dosing of an MDCF for undernourished children as a function of their stage of development and state of disease severity and (iii) evolving policies about complementary feeding practices based on insights about how food components impact the fitness and expressed beneficial functions of growth-promoting elements of a child’s microbiome. Finally, this approach and resulting knowledge base can also serve as a guide for culture initiatives designed to recover candidate probiotic strains, and/or components of synbiotics for repairing gut microbial communities that cannot be resuscitated with food-based interventions alone.

## Methods

### Bacterial genome sequencing and annotation

Monocultures of each isolate were grown overnight at 37 °C in Wilkins-Chalgren Anaerobe Broth (Oxoid Ltd.; catalog number: CM0643) in a Coy Chamber under anaerobic conditions (atmosphere; 75% N_2_, 20% CO_2_ and 5% H_2_) without shaking. Cells were recovered by centrifugation (5000 × g for 10 minutes at 4 °C). High molecular weight genomic DNA was purified (MagAttract® HMW DNA kit, Qiagen) following the manufacturer’s protocol and the amount was quantified (Qubit fluorometer). The sample was passed up and down through a 29-gauge needle 6-8 times and the fragment size distribution was determined (∼30 kbp; TapeStation, Agilent).

Fragmented genomic DNA (400-1000 ng) was prepared for long-read sequencing using a SMRTbell Express Template Prep Kit 2.0 (Pacific Biosciences) adapted to a deep 96-well plate (Fisher Scientific) format. All DNA handling and transfer steps were performed with wide-bore, genomic DNA pipette tips (ART). Barcoded adapters were ligated to A-tailed fragments (overnight incubation at 20 °C) and damaged or partial SMRTbell templates were subsequently removed (SMRTbell Enzyme Cleanup Kit). High molecular weight templates were purified (volume of added undiluted AMPure beads = 0.45 times the volume of the DNA solution). Libraries prepared from different strains were pooled (3-6 libraries/pool). A second round of size selection was then performed; AMPure beads were diluted to a final concentration of 40% (v/v) with SMRTbell elution buffer with the resulting mixture added at 2.2 times the volume of the pooled libraries. DNA was eluted from the AMPure beads with 12 μL of SMRTbell elution buffer. Pooled libraries were quantified (Qubit), their size distribution was assessed (TapeStation) and sequenced [Sequel System, Sequel Binding Kit 3.0 and Sequencing Primer v4 (Pacific Biosystems)]. The resulting reads were demultiplexed and Q20 circular consensus sequencing (CCS) reads were generated (Cromwell workflow configured in SMRT Link software). Genomes were assembled using Flye^37^ (v2.8.1) with HiFi-error set to 0.003, min-overlap set at 2000, and other options set to default. Genome quality was evaluated using CheckM^38^ (v1.1.3) (**Supplementary Table 1a, 4, and 12a**).

We applied Prokka^39^ (v1.14) to identify potential open reading frames (ORF) in each assembled genome. Additional functional annotation of these ORFs using a ‘subsystems’ approach adapted from the SEED genome annotation platform^40^ was performed as described in the accompanying paper^1^. We assigned functions to 9,820 ORFs in 20 isolate genomes using a collection of mcSEED metabolic subsystems that capture the core metabolism of 98 nutrients/metabolites in four major categories (amino acids, vitamins, carbohydrates, and fermentation products) projected over 2,856 annotated human gut bacterial genomes^41–43^. *In silico* reconstructions of selected mcSEED metabolic pathways were based on functional gene annotation and prediction using homology-based methods and genome context analysis. Reconstructions were represented as a binary phenotype matrix (BPM) where, as noted in the main text, assignment of a “1” for amino acids and B vitamins denotes a predicted prototroph and a “0” denotes an auxotroph. For carbohydrates, “1” and “0” refer to a strain’s predicted ability or inability, respectively, to utilize the indicated mono-, di- or oligosaccharide while for fermentation end products, a “1” and “0” indicate a strain’s predicted ability/inability to produce the indicated compound, respectively (see **Supplementary Table 1c, 12d**).

To calculate phylogenetic relationships between five *P. copri* isolates and MAGs Bg0018 and Bg0019, we first used CheckM^38^ (v1.1.3) to extract and align the amino acid sequences of 43 single copy marker genes in each isolate or each of the two MAGs, plus an isolate genome sequence of *Bacteroides thetaiotaomicron* VPI-5482 (PATRIC accession number: 226186.12). Concatenated marker gene sequences were analyzed using fasttree^44^ (v2.1.10) to construct a phylogenetic tree using the Jones-Taylor-Thornton model and CAT evolution rate approximation, followed by tree rescaling using the Gamma20 optimization. The tree was subsequently processed in R using ape^45^ (v5.6-2) to root the tree with the *B. thetaiotaomicron* genome and extract phylogenetic distances between genomes, followed by ggtree^46^ (v3.2.1) for tree plotting.

The similarity between the genomes of these strains and MAGs was quantified by calculating the ANI score with pyani^47^ (ANIm implementation of ANI, v0.2.10). We first calculated ANIm scores for all possible combinations between MAGs and the genomes of cultured bacterial strains and subsequently removed any MAG-strain genome combination with <10% alignment coverage^11^. For the remaining MAGs, a “highly similar” genome in the collection of cultured bacterial strains was defined as having > 94% ANIm score (**Supplementary Table 1b**). We then determined the degree of binary phenotype concordance between each genome in the collection of cultured bacterial strains and its “highly similar” MAG. A binary phenotype concordance score was calculated by dividing the number of binary phenotypes^1^ shared between a cultured strain’s genome and a MAG by the total number of binary phenotypes annotated in the strain and MAG. A ‘Representative MAG’ for each genome was defined as having a binary phenotype concordance score >90% (**Supplementary Table 1c**).

PULs were predicted based on the method described in ref. 48 and displayed with the PULDB interface^49^. PULs were placed into three categories: (i) ‘functionally conserved’ (PULs containing shared ORFs encoding the same CAZymes and SusC/SusD proteins in the same organization in their respective genomes with ≥90% amino acid identity between proteins); (ii) ‘structurally distinct’ (PULs present in respective genomes but where one or more CAZymes or one or both SusC/SusD proteins are missing or fragmented in a way likely to impact function, or where extra PUL elements are present), and (iii) ‘not conserved’ (PULs present in respective genomes but with mutations likely to completely compromise function, or no PUL identified).

### Colonization and husbandry

Gnotobiotic mouse experiments were performed using protocols approved by the Washington University Animal Studies Committee. Germ-free C57BL/6J mice were maintained in plastic flexible film isolators (Class Biologically Clean Ltd) at 23 °C under a strict 12-hour light cycle (lights on a 0600h). Autoclaved paper ‘shepherd shacks’ were kept in each cage to facilitate natural nesting behaviors and provide environmental enrichment.

A weaning diet containing MDCF-2 was formulated as described in the main text. Ingredients represented in the different diet modules were combined and the mixture was dried, pelleted, and sterilized by gamma irradiation (30-50 Kgy). Sterility was confirmed by culturing the pellets in LYBHI medium^50^ and in Wilkins-Chalgren Anaerobe Broth under aerobic and anaerobic conditions for 7 days at 37 °C followed by plating on LYBHI- and blood-agar plates. Nutritional analysis of each irradiated diet was performed by Nestlé Purina Analytical Laboratories (St. Louis, MO) (**Supplementary Table 2c**).

Pregnant C57BL/6J mice originating from trio matings were given *ad libitum* access to an autoclaved breeder chow (Purina Mills; Lab Diet 5021) throughout their pregnancy and until postpartum day 2. Key points about the experimental design of the gnotobiotic mouse experiments described in **Fig. 1b, Extended Data Fig. 7a, Extended Data Fig. 8a**, and **Fig. 4a** are; (i) all bacterial strains were cultured in Wilkins-Chalgren Anaerobe Broth (except for *F. prausnitzii* which was cultured in LYBHI medium) and were harvested after overnight growth at 37 °C (**Supplementary Table 1a**), (ii) all gavage mixtures contained equivalent amounts (by OD_600_) of their constituent bacterial strains except for *F. prausnitzii* which was concentrated 100-fold before preparing the gavage mixture, (iii) each bacterial consortium was administered to the postpartum dams in a volume of 200 μL using an oral gavage needle (Cadence Science; catalog number: 7901), (iv) the number of dams and pups per treatment group (two dams and 7-8 pups/treatment group for the experiment described in **Fig. 1b**; four dams and 18-19 pups/treatment group for the experiment outlined in **Extended Data Fig. 7a**; two dams and 13 pups/treatment group for the experiment illustrated in **Fig. 4a**; two dams and 12 pups/treatment group for the experiment shown in **Extended Data Fig. 8a**), (v) half of the bedding was replaced with fresh bedding in each cage each day from postpartum day 1 to 14, after which time bedding was changed every 7 days, (vi) diets were provided to mothers as well as to their weaning and post-weaning pups *ad libitum*, and (vii) biospecimens collected from mice when they were euthanized (without prior fasting) were snap frozen in liquid nitrogen and stored at −80 °C before use.

Pups were weighed on P23, P35, and P53, and normalized to the weight on P23. A linear mixed-effects model was used to evaluate the effect of different microbial communities on normalized mouse weight gain:

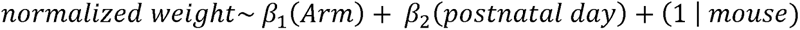

### Defining the absolute abundances of bacterial strains in ileal, cecal and fecal communities

The absolute abundances of bacterial strains were determined in the microbiota using previously described methods with minor modifications^51–52^. In brief, 3.3×10^6^ cells of *Alicyclobacillus acidiphilus* DSM 14558 and 1.49×10^7^ cells of *Agrobacterium radiobacter* DSM 30147 (ref. 51) were added to each weighed frozen sample prior to DNA isolation and preparation of barcoded libraries for shotgun sequencing. Sequencing was performed using an Illumina NextSeq instrument. Bacterial abundances were determined by assigning reads to each bacterial genome, followed by a normalization for genome uniqueness in the context of a given community^52^. The resulting count table was imported into R^53^ (v4.0.4). We calculated the absolute abundance of a given strain *i* in sample *j* in reference to the spike-in

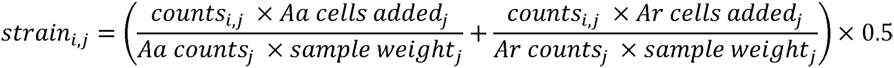

The statistical significance of observed differences in the abundance of a given strain between experimental groups in ileal, cecal, and fecal samples was determined by using the Kruskall-Wallis test followed by Dunn’s test for each pairwise comparison among the three arms in the experiment described in **Fig. 1** or the Mann-Whitney U test between the two arms in experiments described in **Extended Data Fig. 1b, Extended Data Fig. 7**, and **Fig. 4**. The statistical significance of differences in the composition of communities in the three arms described in **Fig. 1g** was determined using PERMANOVA^54^ on sample projections onto principal components calculated from the log_10_ absolute abundance profiles of the 16 organisms that were not *B. infantis* or *Prevotella* species.

For the experiment described in **Fig. 1**, 96 fecal samples were sequenced [2.2×10^6^±1.2×10^5^ unidirectional 75 nt reads/sample (mean ± SD)], along with 20 cecal samples [1.5×10^6^±4.2×10^5^ unidirectional 75 nt reads/sample] and 20 ileal samples [1.5×10^6^±9.1×10^4^ unidirectional 75 nt reads/sample] (**Supplementary Table 3a**). For the experiment described in **Extended Data Fig. 1b**, 37 fecal samples were sequenced [5.8×10^6^±1.6×10^6^ unidirectional 75 nt reads/sample] (**Supplementary Table 5a)**], while for the experiment described in **Extended Data Fig. 7a**, 37 cecal samples were sequenced [1.3×10^6^ ± 1.3×10^5^ unidirectional 75 nt reads/sample (**Supplementary Table 10a**)]. For the experiment described in **Fig. 4b**, 26 cecal samples were sequenced [2.9×10^6^ ± 5.6×10^6^ 150nt paired-end reads/samples (**Supplementary Table 13a**)].

The statistical significance of observed differences in the abundance of a given strain across different treatment groups and time was tested using a linear mixed effects model within the R packages lme4^55^ (v1.1-27) and lmerTest^56^ (v3.1-3). The change in *P. copri* absolute abundance in fecal samples during the course of the experiment was determined by a linear mixed-effects model:

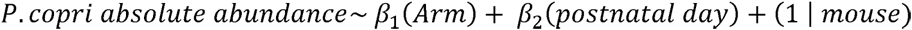

### Microbial RNA-Seq

RNA was isolated^5^ from cecal contents collected at the end of the experiment. cDNA libraries were generated from isolated RNA samples using the ‘Total RNA Prep with Ribo-Zero Plus’ kit (Illumina). Barcoded libraries were sequenced [Illumina NovaSeq instrument]. For the experiment described in **Fig. 1b**, cDNA recovered from 20 different samples of cecal contents were each sequenced to a depth of 7.8×10^7^ ± 9.6×10^6^ bidirectional 150 nt reads (mean±SD) (**Supplementary Table 7a**). For the experiment shown in **Fig. 4b**, DNA from 26 different samples of cecal contents were each sequenced to a depth of 6.5×10^7^ ± 2.1×10^7^ bidirectional 150 nt reads (**Supplementary Table 15a**). Raw reads were trimmed by using TrimGalore^57^ (v0.6.4). Trimmed reads longer than 100 bp were mapped to reference genomes with kallisto^58^ (v0.43.0).

To analyze expression of genes from the *P. copri* and *P. stercorea* PULs described in **Extended Data Fig. 3a-b** and **Fig. 4e**, transcripts per million (TPM) values were by mapping reads, using kallisto, to their genomes (see **Fig. 1b**, **Fig. 4b**). TPM normalized expression was used to control for differences in library depth and gene length. For **Extended Data Fig. 3a-b** and **Fig. 4e**, log_2_ TPM with a pseudocount of 1 were visualized for all predicted PUL genes using the seaborn^59^ (v0.12.1) Python package, splitting the set to show individual PULs in color on the right side of each violin plot against the remainder of the PUL genes shown in gray on the left side of each violin plot. Adjusted GSEA *P*-values were calculated with fgsea^60^ (v1.20.0), ranking genes by their mean log_2_ TPM across the *Prevotella*-colonized samples in a given arm. Each PUL comprised a gene set against the background of all PUL genes from a given isolate, with minimum and maximum gene set sizes of 5 and 50 genes, respectively (**Supplementary Table 7b, Supplementary Table 15b**).

For the analysis described in **Fig. 4e**, modifications were made to account for the high fraction of genes with 100% nucleotide identity between *P. copri* BgD5_2 and BgF5_2. The kallisto index was generated to represent the set of unique genes between the two isolates by including the entire *P. copri* BgF5_2 gene set and only the subset of *P. copri* BgD5_2 genes that did not share identical sequences in *P. copri* BgF5_2 (61 out of 3,066 genes). TPMs were filtered down to the set of unique genes in the *P. copri* BgD5_2 and BgF5_2 PULs [n=201 unique genes, consisting of 195 genes with identical nucleotide sequences between the two isolates and 3 unique genes from each isolate (**Supplementary Table 12c**)].

For abundance-normalized differential expression analysis of microbial transcripts described in **Fig. 2d**, paired metagenomic and meta-transcriptomic kallisto pseudocounts were generated by mapping reads from the cecal DNA and cDNA libraries described above to the specific set of bacteria administered in the arm from which the sample was derived. To prevent skewing of library size normalization, the resulting counts matrix then underwent filtering of counts for rRNA loci predicted by Prokka^39^. *MTXmodel*^18^ was run using the paired metagenomic and meta-transcriptomic counts tables for each organism to achieve ‘within-taxon-sum-scaling.’^61^ Each pairwise comparison of arms was run using the respective sets of samples with ‘arm’ as the single fixed effect in the generalized linear model design. Transcripts with statistically significant differences in their expression after normalizing for absolute abundance were identified [q-value (adjusted *P*-value) < 0.1] after multiple hypothesis correction was applied to the entire set of transcripts from a given organism. GSEA was performed for each organism with fgsea^60^, ranking all genes surviving zero-filtering from *MTXmodel* differential expression testing by their estimated log fold-difference. Each mcSEED metabolic pathway in each organism was used as a gene set against the background of all genes tested for differential expression, with minimum and maximum gene set sizes of 5 and 50 genes, respectively.

### Histomorphometric analysis of villus height and crypt depth

Jejunal and ileal segments were fixed in formalin, embedded vertically in paraffin, 5 μm-thick sections were prepared and the sections were stained with hematoxylin and eosin. Slides were scanned (NanoZoomer instrument, Hamamatsu). For each animal, 10 well-oriented crypt-villus units were selected from each intestinal segment for measurement of villus height and crypt depth using QuPath^62^ (v0.3.2). Measurements were performed with the investigator blinded with respect to colonization group. A two-tailed Mann-Whitney U test was applied to the resulting datasets.

### Single nucleus (sn) RNA-seq

Jejunal segments of 1.5 cm in length were collected from mice and snap frozen in liquid nitrogen [n=4 animals/treatment group (2 males and 2 females); 2 treatment groups in total]. The method for extracting nuclei was adapted from a previously described protocol for the pancreas^63^. Briefly, tissues were thawed and minced in lysis buffer [25mM citric acid, 0.25M sucrose, 0.1% NP-40, and 1× protease inhibitor (Roche)]. Nuclei were released from cells using a pestle douncer (Wheaton), washed 3 times with buffer [25mM citric acid, 0.25M sucrose, and 1× protease inhibitor], and filtered successively through 100μm, 70μm, 40μm, 20μm and finally 5μm diameter strainers (pluriSelect) to obtain single nuclei in resuspension buffer [25mM KCl, 3mM MgCl_2_, 50mM Tris, 1mM DTT, 0.4U/μL rNase inhibitor (Sigma) and 0.4U/μL Superase inhibitor (ThermoFisher)]. Approximately 10,000 nuclei per sample were subjected to gel bead-in-emulsion (GEM) generation, reverse transcription and construction of libraries for sequencing according to the protocol provided in the 3’ gene expression v3.1 kit manual (10X Genomics). Libraries were balanced, pooled and sequenced [Illumina NovaSeq S4; 3.23×10^8^ ± 1.39×10^7^ paired-end 150 nt reads/nucleus (mean±SD) from jejunal samples]. Read alignment, feature-barcode matrices and quality controls were processed by using the 10X Genomics CellRanger^64^ 5.0 pipeline with the flag ‘-include-introns’ to ensure that reads would be allowed to map to intronic regions of the mouse reference genome (GRCm38/mm10). Nuclei with over 2.5% reads from mitochondria-encoded genes or ribosomal protein genes were filtered out.

#### Analysis of snRNA-seq datasets

Sample integration, count normalization, cell clustering and marker gene identification was performed using Seurat 4.0^22^. Briefly, filtered feature-barcode matrices outputted from CellRanger were imported as a Seurat object using *CreateSeuratObject* (min.cells = 5, min.features = 200). Each sample was normalized using *SCTransform*^65,66^ and integrated using *SelectIntegrationFeatures*, *PrepSCTIntegration*, *FindIntegrationAnchors*, and *IntegrateData* from the Seurat software package. The integrated dataset, incorporating nuclei from all samples, was subjected to unsupervised clustering using *FindNeighbors* (dimensions = 1:30) and *FindClusters* (resolution = 1) from the Seurat package, which executes a shared nearest-neighbor graph clustering algorithm to identify putative cell clusters. Cell type assignment was performed manually based on expression of reported markers^34,67,68^.

Cross-condition differential gene expression analysis was performed based on a “pseudobulk” strategy: for each cell cluster, gene counts were aggregated to obtain sample-level counts; each pseudo-bulked sample served as an input for edgeR-based differential gene expression analysis^19,20^.

For *NicheNet*-based analysis^21^ (v1.1.0), all clusters in our snRNA-seq dataset were used as senders for crypt stem cells, proliferating TA/stem cells, villus base enterocytes, mid-villus enterocytes and villus tip enterocytes, plus goblet cells. We used the *nichenet_seuratobj_aggregate* (assay_oi = “RNA”) function with its default settings to incorporate differential gene expression information from Seurat into our *NicheNet* analysis and to select bona fide ligand-receptor interactions.

*Compass*-based *in silico* metabolic flux analysis^24^ (v0.9.10.2) was performed using transcripts from each of six epithelial cell clusters (crypt stem cells, proliferating TA cells, villus-base, mid-villus and villus tip enterocytes and goblet cells). The reaction scores calculated by *Compass* were filtered based on (i) the confidence levels of the Recon2 reactions and (ii) the completeness of information for Recon2 reaction annotations. Only Recon2 reactions that are supported by biochemical evidence (defined by Recon2 as having a confidence level of 4; ref. 25) and that have complete enzymatic information for the reaction were advanced to the follow-on analysis (yield: 2,075 pass filter reactions in 83 Recon2 subsystems).

We subsequently calculated a “metabolic flux difference” to determine whether the presence or absence of *P. copri* affected *Compass-*based predictions of metabolic activities at the Recon2 reaction level in the six cell clusters. The “net reaction score” was calculated as follows

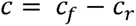

Where *C_f_* denotes the *Compass* score for a given reaction in the “forward” direction, and, if the biochemical reaction is reversible, *C_r_* denotes the score for the “reverse” reaction.

A Wilcoxon Rank Sum test was used to test significance of the net reaction score between the two treatment groups. *P* values from the Wilcoxon Rank Sum tests were adjusted for multiple comparisons with the Benjamini-Hochberg method.

Cohen’s *d* can be used to show the effect size of *c_f_* or *c_r_* for each reaction between two groups^24^ (in our case mice harboring communities with and without *P. copri*). Briefly, Cohen’s *d* of two groups, *j* and *k*, was calculated based on the following two equations^69^ where *n, s,* and *a* represent the number, the variance, and the mean of the observations (in our case, the net reaction scores).

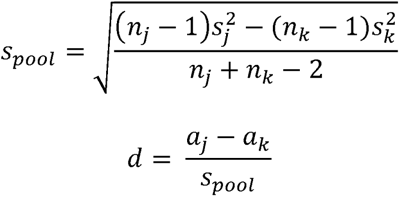

If both *a_j_* and *a_k_* are non-negative numbers, a positive Cohen’s *d* indicates that the mean of group *j* is greater than that of group *k,* whereas a negative Cohen’s *d* means the mean of group *j* is smaller in that comparison. The magnitude of Cohen’s represents the effect size and is correlated with the difference between the means of the two groups. Because the mean of the net subsystem scores as well as the net reaction scores could be negative, we made the following adjustments to Cohen’s *d* in order to preserve the concordance of sign and the order of group means. The adjusted Cohen’s *d* represents the metabolic flux difference *m,* and is defined as:

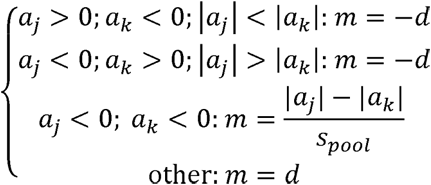

*scCODA*^27^ (v0.1.8) is a Bayesian probabilistic model for detecting ‘statistically credible differences’ in the proportional representation of cell clusters, identified from snRNA-seq datasets, between different treatment conditions^27^. This method accounts for two main challenges when analyzing snRNA-seq data: (i) low sample number and (ii) the compositionality of the dataset (an increase in the proportional representation of a specific cell cluster will inevitably lead to decreases in the proportional representation of all other cell clusters). Therefore, applying univariate statistical tests, such as a t-test, without accounting for this inherent negative correlation bias will result in reported false positives. *scCODA* uses a Bayesian generalized linear multivariate regression model to describe the ‘effect’ of treatment groups on the proportional representation of each cell cluster; Hamiltonian Monte Carlo sampling is employed to calculate the posterior inclusion probability of including the effect of treatment in the model. The type I error (false discovery) is derived from the posterior inclusion probability for each effect. The set of “statistically credible effects” is the largest set of effects that can be chosen without exceeding a user-defined false discovery threshold α (α*=*0.05 by default). We applied *scCODA* using default parameters, including choice of prior probability in the Bayesian model and the setting for Hamiltonian Monte Carlo sampling. The enteroendocrine cell cluster was used as the reference cluster in accordance with recommendations from the creators of *scCODA* to choose a cell cluster that has consistent proportional representation across samples.

### Mass spectrometry

#### UHPLC-QqQ-MS of cecal glycosidic linkages and GC-MS of short-chain fatty acids

Ultra-high performance liquid chromatography-triple quadrupole mass spectrometric (UHPLC-QqQ-MS) quantification of glycosidic linkages and monosaccharides present in cecal glycans was performed using methods described in the accompanying study^1^. Levels of short-chain fatty acid levels in cecal contents were measured by GC-MS using a procedure outlined in ref. 5.

#### LC-MS of acylcarnitines, amino acids, and biogenic amines in host tissues

Acylcarnitines were measured in jejunum, colon, liver, gastrocnemius, quadriceps, and heart muscle, plus plasma according to ref. 70, while 20 amino acids plus 19 biogenic amines were quantified in jejunum, liver, and muscle according to methods detailed in ref. 71. Plasma levels of non-esterified fatty acids were measured using a UniCel DxC600 clinical analyzer (Beckman Coulter).

#### Targeted mass spectrometry of cecal amino acids and B vitamins

Methods for targeted LC-QqQ-MS of amino acids and B vitamins were adapted from a previous publication^72^. Cecal samples were extracted with ice-cold methanol, and a 200μL aliquot was dried (vacuum centrifugation; LabConco CentriVap) and reconstituted with 200μL of a solution containing 80% methanol in water. A 2μL aliquot of extracted metabolites was then injected into an Agilent 1290 Infinity II UHPLC system coupled with an Agilent 6470 QqQ-MS operated in positive ion dynamic multiple reaction monitor mode (dMRM). The native metabolites were separated on HILIC column (ACQUITY BEH Amide, 2.1 x 150 mm, 1.7 μm particle size, Waters) using a 20-minute binary gradient with constant flow rate of 0.4 mL/minute. The mobile phases were composed of 10mM ammonium formate buffer in water with 0.125% formic acid (Phase A) and 10mM ammonium formate in 95% acetonitrile/H_2_O (v/v) with 0.125% formic acid (Phase B). The binary gradient was as follows: 0-8 minutes, 91-90% B; 8-14 minutes, 90-70% B; 15-15.1 minutes, 70-91% B; 15.1-20 minutes, 91% B. A pool of 20 amino acids and 7 B vitamins standards with known concentrations (amino acid pool: 0.1ng/mL-100μg/mL; B vitamin pool: 0.01ng/mL-10μg/mL) was injected along with the samples as an external calibration curve for absolute quantification.

## Acknowledgments

We thank David O’Donnell and Maria Karlsson for their invaluable assistance with mouse husbandry, Martin Meier for his essential role in generating shotgun sequencing and microbial RNA-Seq datasets, Janaki Guruge for her help with culturing and maintaining bacterial strains, plus Su Deng and Justin Serugo for oversight of biospecimen archives. This work was supported by grants from the NIH (DK30292, T32 HG000045), the Centene-Washington University Personalized Medicine Initiative and the Bill & Melinda Gates Foundation. Y.W. is the recipient of a career development award from the NIH (K01 DK134840).

## Author Contributions

H.-W.C., E.M.L., Y.W., C.Z. and J.I.G. designed the gnotobiotic mouse experiments, which were performed by H.-W.C., E.M.L. and C.Z.. Mouse diets were formulated by H.-W.C. with assistance from M.J.B. and were based on the results of human diet surveys provided by I.M, S.D., M.M. and T.A.. Bacterial genomes were sequenced and assembled by D.M.W., H.-W.C. and M.C. with S.H., D.A.R., A.A.A., N.T., B.H. and A.L.O. contributing to genome annotation. Shotgun sequencing and microbial RNA-Seq datasets were generated by H.-W.C. and E.M.L., and analyzed together with M.C.H., H.M.L., and D.M.W.. Mass spectrometry-based metabolomic studies were conducted by J.C., O.I., M.J.M. and C.B.N.. Glycan analyses were performed by J.J.C., G.C., Y.C., N.P.B. and C.B.L.. Morphometric analysis was conducted by Y.W.. snRNA-Seq datasets were generated and analyzed by H.-W.C., Y.W., K.M.P., R.Y.C., C.K. and M.C.. H.-W.C., E.M.L., Y.W. and J.I.G. wrote this manuscript with invaluable assistance from co-authors.

## Materials & Correspondence

Correspondence and material requests should be sent to Jeffrey Gordon

---

**Extended Data Fig. 1.**
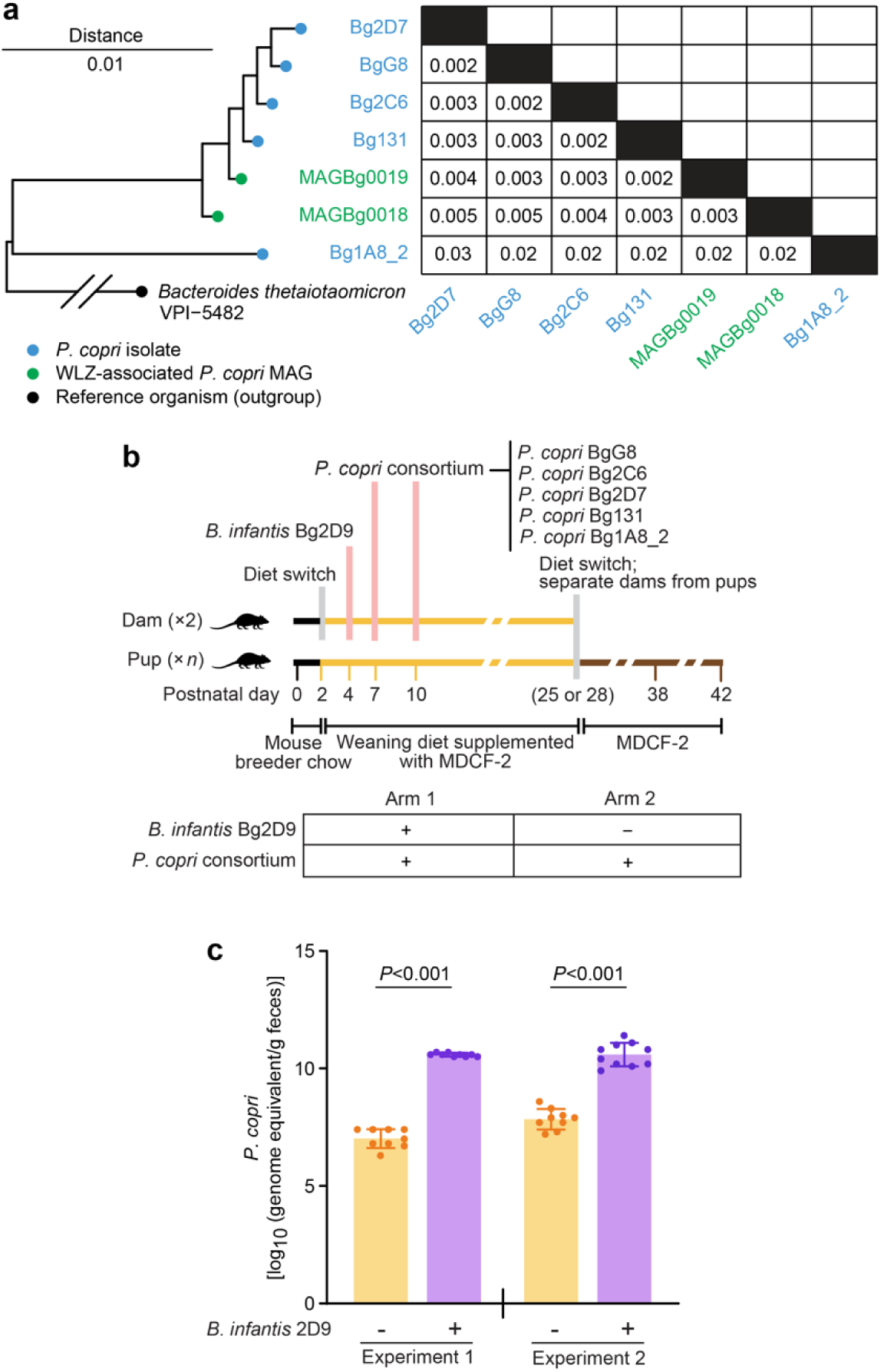
Determining the relationship between *P. copri* colonization efficiency and pre-colonization with *B. longum* subsp. *infantis*. (**a)** Phylogenetic tree of cultured *P. copri* isolates used in the mouse studies described and the two MAGs positively associated with WLZ in the randomized controlled human study. The phylogenetic distance between each pair of comparisons is shown in the matrix. **(b)** Experimental design. Mice were weaned at P28 and P25 for experiments 1 and 2, respectively. **(c**) Total absolute abundances of *P. copri* strains in fecal samples collected from mice at P42. Mean values ± SD are shown. Each dot represents a separate animal. *P*-values were calculated using a Mann-Whitney U test.

**Extended Data Fig. 2.**
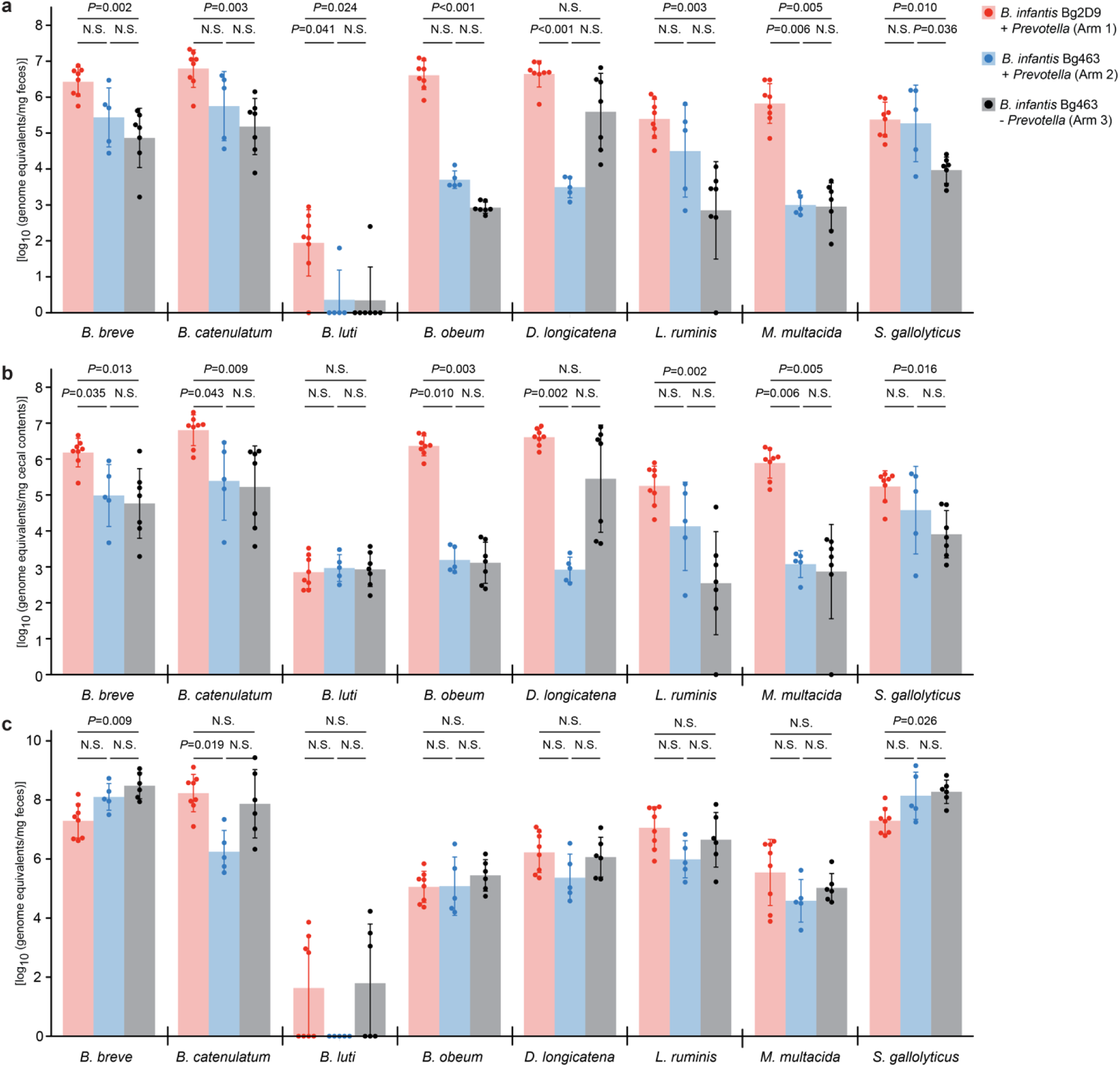
Absolute abundances of other bacterial species in the defined community across different time points and locations. (**a)** Absolute abundances of organisms which were significantly higher with either *B. infantis* Bg2D9 (Arm 1 vs. Arm 2), or with the combination of *B. infantis* Bg2D9 and *Prevotella* (Arm 1 vs. Arm 3) in fecal samples collected at P53. (**b, c**) Absolute abundances of the same organisms in cecal contents collected at P53 (panel b) and in fecal samples collected before weaning at P21 (panel c). Mean values ± SD are shown. Each dot represents an individual animal. *P*-values were calculated by the Kruskal-Wallis test followed by post-hoc Dunn’s test with Bonferroni correction. N.S., not significant (*P>*0.05).

**Extended Data Fig. 3.**
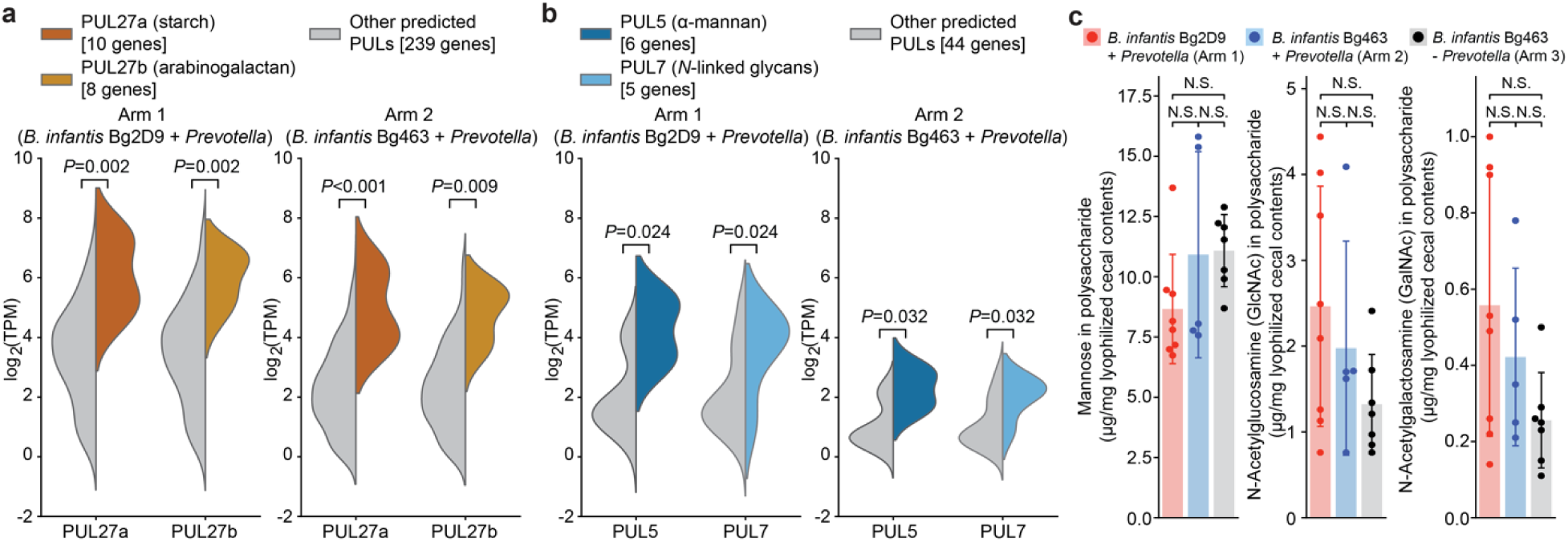
Expression of *P. copri* and *P. stercorea* PULs and targeted mass spectrometric analysis of their predicted targets. (**a,b)** GSEA of expression of PULs shared by *P. copri* Bg131 (panel a) and *P. stercorea* (panel b) in the two *Prevotella-*containing arms of the experiment described in Fig. 1. Adjusted *P*-values were calculated using GSEA ranking genes by their mean log_2_ TPM across the *Prevotella*-colonized samples in Arms 1 and 2, with each PUL comprising a gene set against the background of all predicted PUL genes. Violin plots show the log_2_ TPM of all genes assigned to any of the PULs in each isolate; plots are split to show the indicated PUL. (**c**) UHPLC-QqQ-MS-based quantitation of levels of total mannose, N-acetylglucosamine, and N-acetylgalactosamine in cecal glycans collected at P53. Mean values ± SD are shown. Each dot represents an individual animal. *P*-values were calculated by the Kruskal-Wallis test followed by post-hoc Dunn’s test with Bonferroni correction.

**Extended Data Fig. 4.**
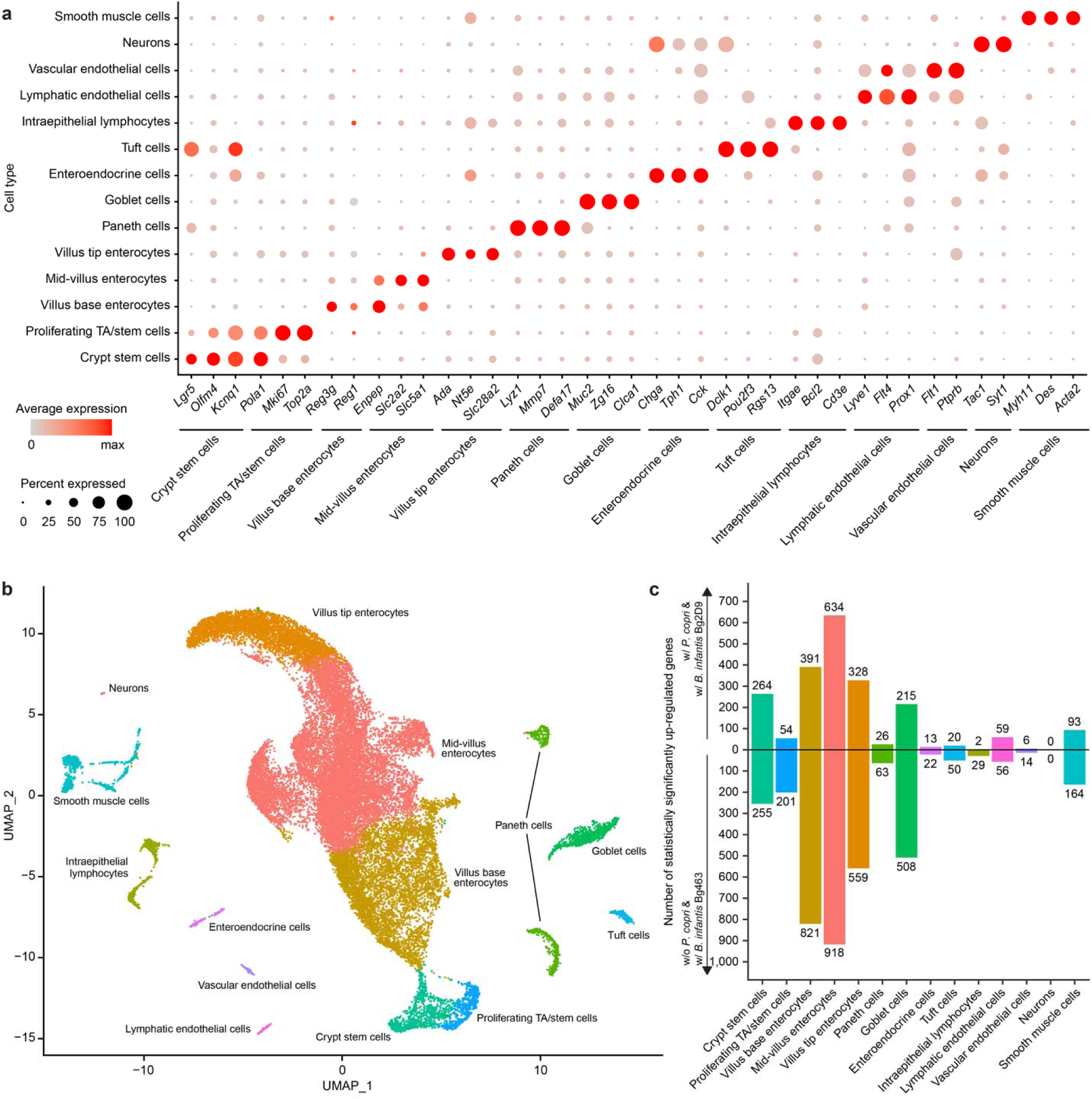
snRNA-Seq analysis of differential intestinal gene expression in mice colonized with bacterial consortia with or without *P. copri*. Jejunal tissue samples collected from ‘w/ *P. copri* & w/ *B. infantis* Bg2D9’ and ‘w/o *P. copri* & w/ *B. infantis* Bg463’ groups at the end of the experiment described in Fig. 1 were analyzed. (**a**) Dot plot of marker gene expression across epithelial cell types. The average expression level and percentage of nuclei that express a given gene within a cell type are indicated by dot color and size, respectively. (**b**) Integrated UMAP plot for all jejunal nuclei isolated from animals in both arms (n=4 mice/arm). (**c**) The number and directionality of statistically significant differentially expressed genes in each cell cluster.

**Extended Data Fig. 5.**
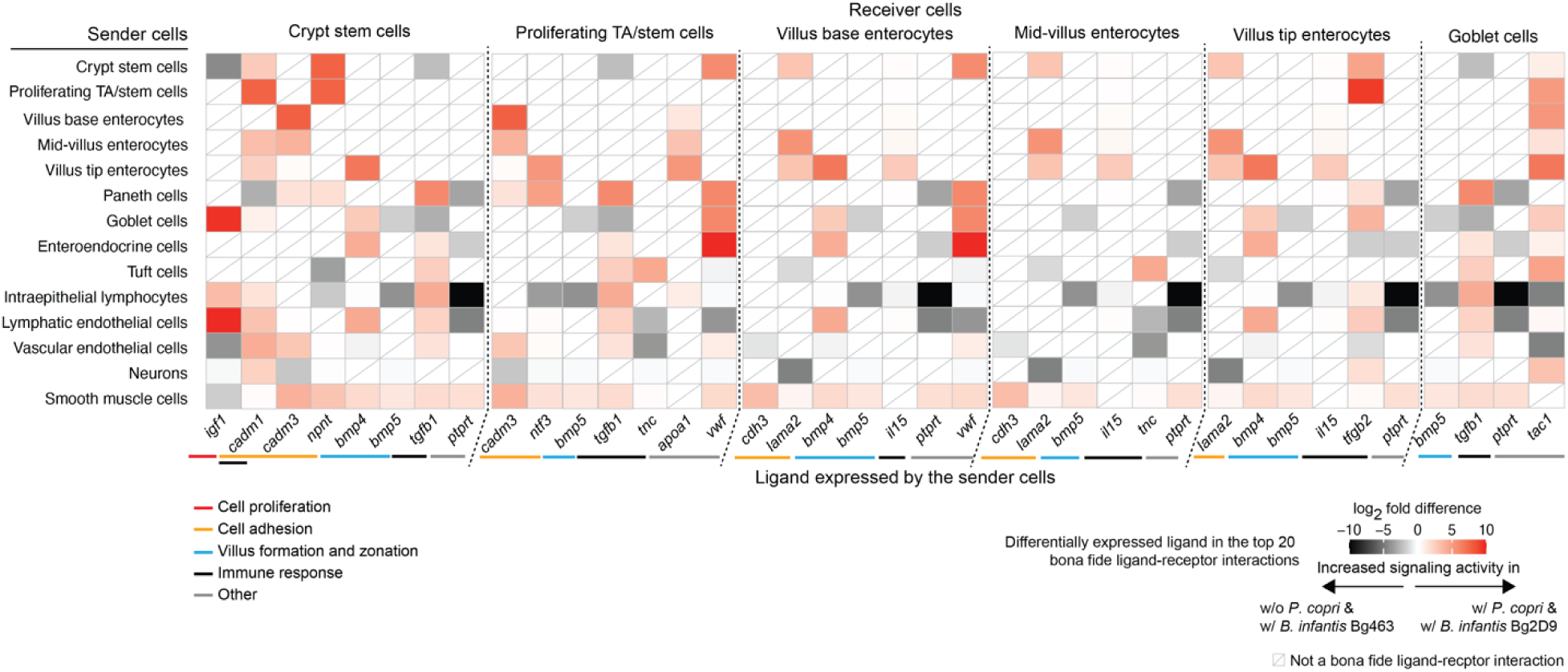
*NicheNet*-based analysis of the effects of *P. copri* colonization on cell-cell signaling activities. Each row represents different sender cell clusters. Each column represents ligands expressed by these sender cells. Cells are colored based on the log_2_-fold difference in expression of ligands in the sender cell clusters between mice in ‘w/ *P. copri* & w/ *B. infantis* Bg2D9’ and ‘w/o *P. copri* & w/ *B. infantis* Bg463’ groups from the experiment described in Fig. 1. Ligands (columns) are grouped based on receiver cell clusters and the indicated functions of downstream signaling pathways in these receiver cells.

**Extended Data Fig. 6.**
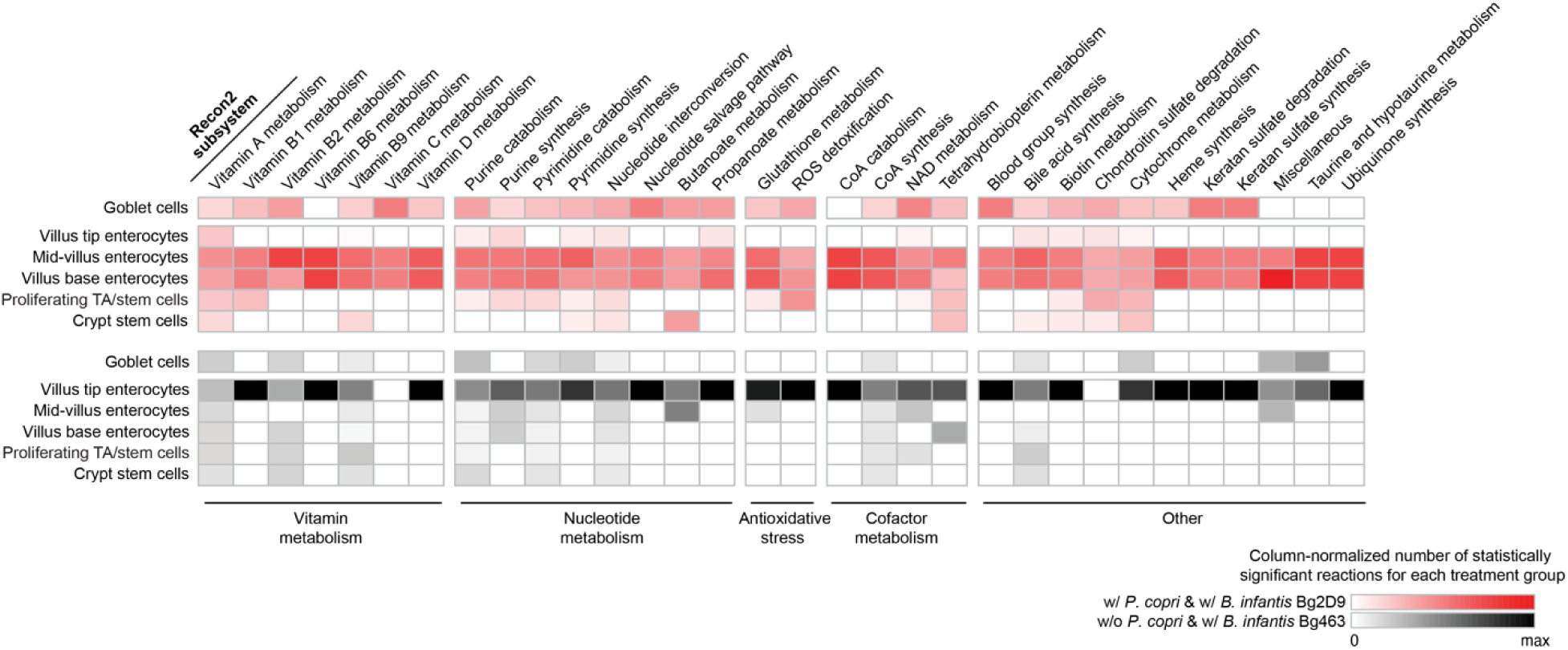
Normalized number of Recon2 reactions in Recon2 subsystems predicted to have statistically significant differences in their activities between mice in the ‘w/ *P. copri* & w/ *B. infantis* Bg2D9’ and ‘w/o *P. copri* & w/ *B. infantis* Bg463’ groups. See legend to Fig. 3b, which shows other affected subsystems, for details.

**Extended Data Fig. 7.**
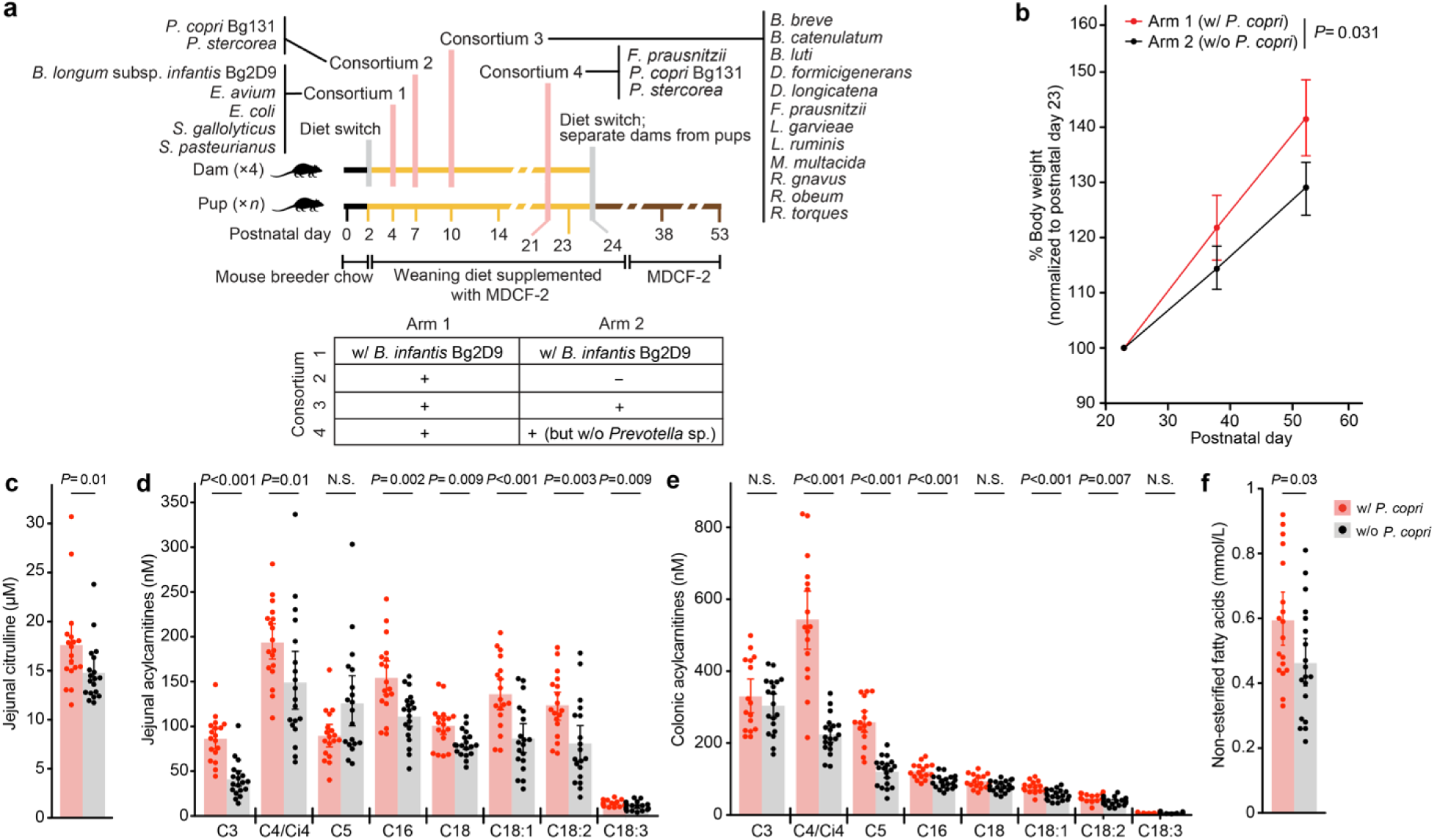
Validating the effects of *P. copri* colonization on postnatal weight gain and host metabolism in gnotobiotic mother-pup dyads. (**a**) Study design (n=4 dams and 18-19 offspring/ treatment arm). (**b**) Body weights of the offspring of dams, normalized to postnatal day 23 [linear mixed effects model (see *Methods*)]. (**c-e**) Targeted mass spectrometric analysis of jejunal citrulline (panel c) and acylcarnitine levels (panel d), plus colonic acylcarnitine levels (panel e). (**f**) Plasma levels of non-esterified fatty acids. Each dot represents a single animal. Mean values ± SD are shown for panels b-f. *P*-values were calculated from the linear mixed effect model (panel b) or Mann-Whitney U test (panels c-f). N.S., *P*-value > 0.05.

**Extended Data Fig. 8.**
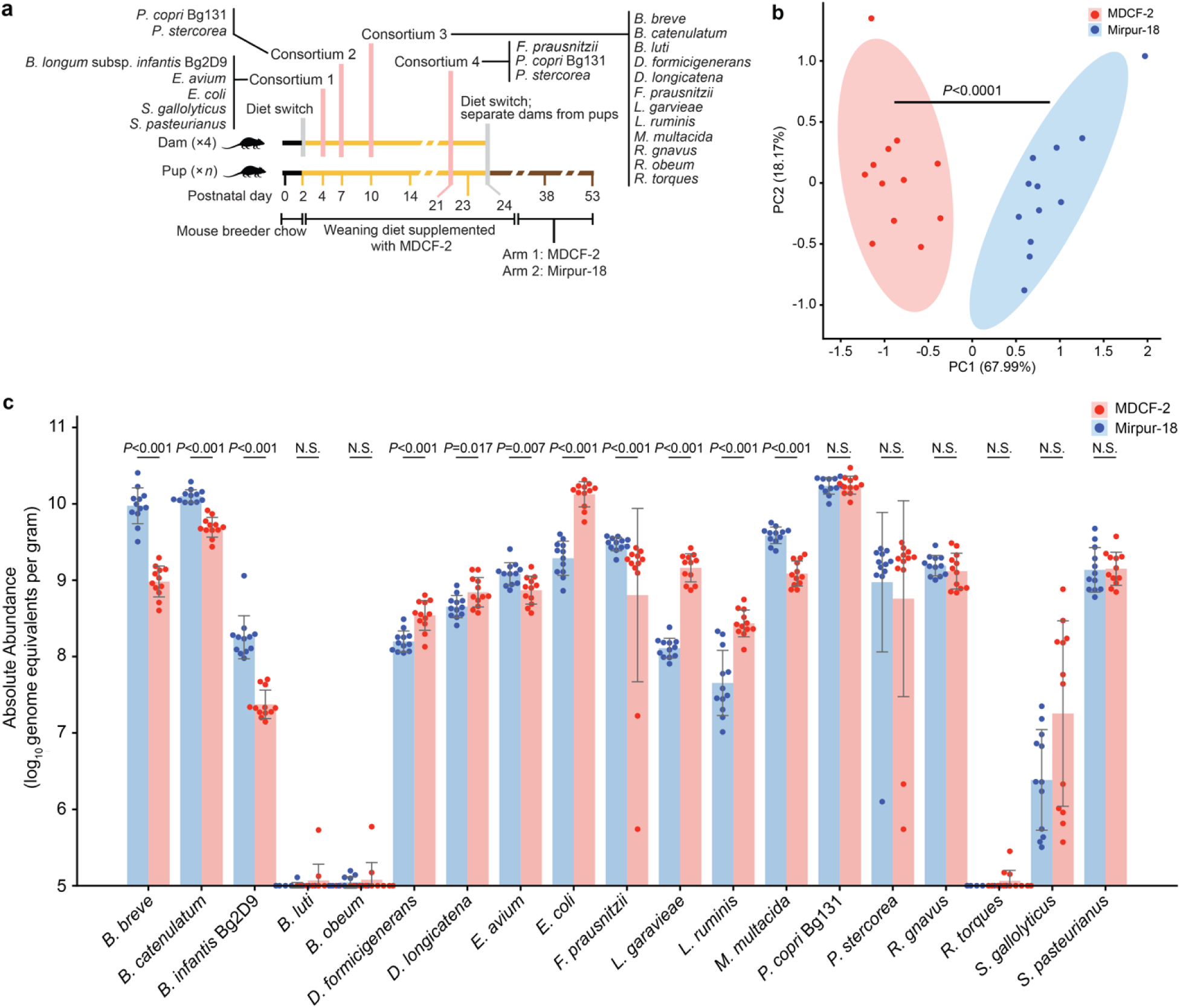
Evaluating the effect of diet on the defined community in gnotobiotic dam-pup dyads. (a) Experimental design (n=2 dams and 12 offspring/diet treatment). **(b)** Principal component analysis showing the significant differences in community structure in the cecums of mice euthanized on P53 (*P*<0.0001; PERMANOVA). Ellipses represent 95% confidence intervals. **(c)** Absolute abundances of the defined community members in cecal contents at P53; *P*-values were calculated by the Mann-Whitney U test. N.S., *P*-value > 0.05. Mean values ± SD are shown. Each dot represents an individual animal.

**Extended Data Fig. 9.**
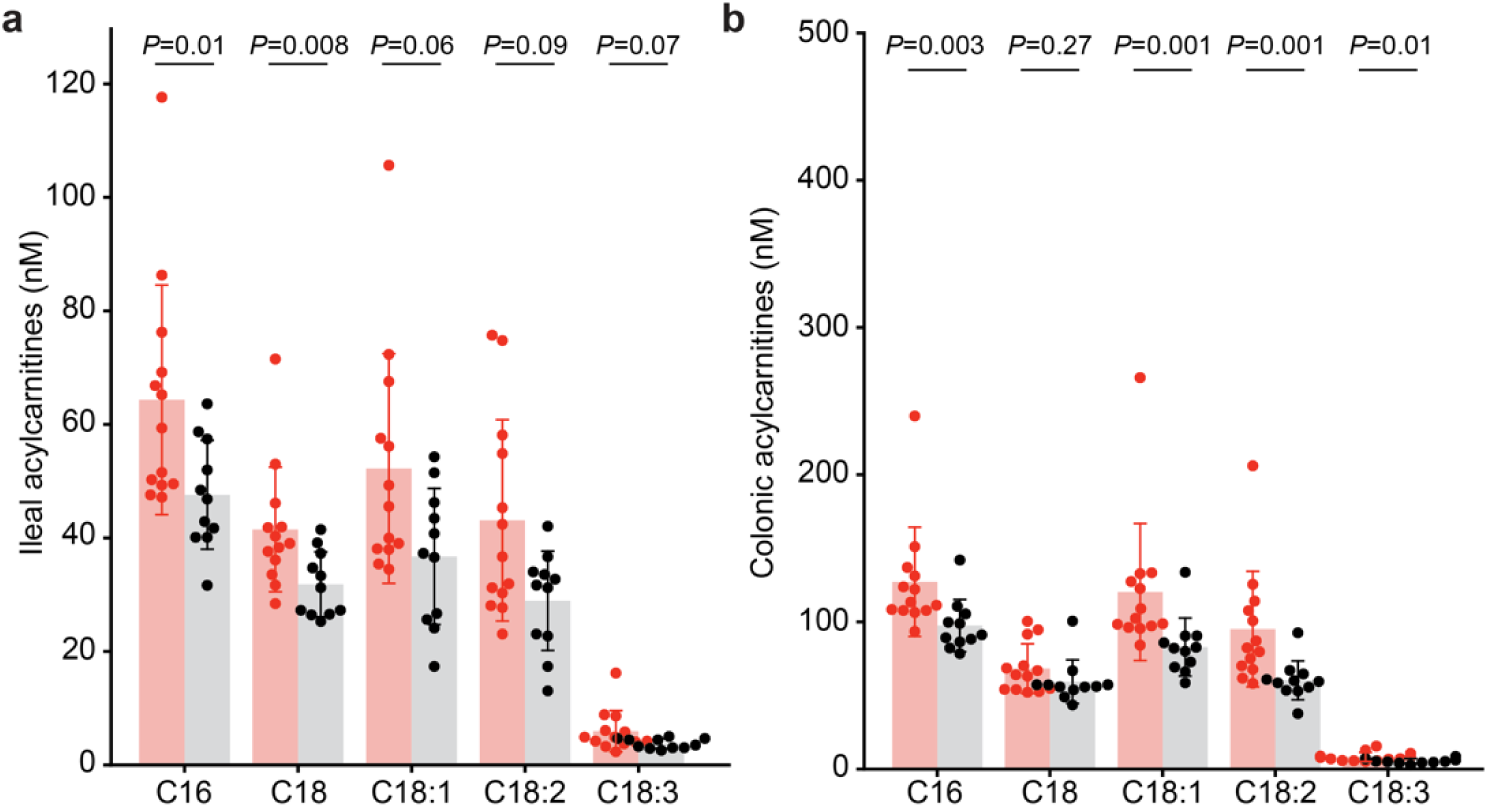
LC-MS of ileal and colonic acylcarnitines in gnotobiotic mice colonized with *P. copri* BgD5_2 and BgF5_2. (a) LC-MS of ileal acylcarnitines corresponding to soybean oil lipids. (b) LC-MS of colonic acylcarnitines corresponding to soybean oil lipids. Each dot represents an individual animal. Mean values ± SD are shown. *P-*values were calculated using a Mann-Whitney U test.

## Supplementary Information

### Data Availability

Microbial community and bacterial strain genome shotgun sequencing datasets plus microbial RNA-Seq and snRNA-Seq datasets have been deposited in the European Nucleotide Archive under study accession PRJEB47807. UHPLC-QqQ-MS datasets are available in Glycopost (ID GPST000245).

### Biological Materials

Human specimens and bacterial strains cultured from fecal samples collected from Bangladeshi children are the property of icddr,b. Material Transfer Agreements exist between icddr,b, and Washington University for the use of these samples. Requests for materials should be made to J.I.G.

### Computer Code

All software used was from publicly available sources.

## Supplementary Tables

**Supplementary Table 1. Bacterial strains used in the defined community gnotobiotic mouse experiments**. (**a**) Quality metrics for assembled bacterial genomes. (**b**) Comparison between the genomes of cultured bacterial strains and MAGs. (**c**) Binary phenotype predictions for cultured bacterial strains in the 20-member consortium. (**d**) Comparison of the representation of metabolic pathways in the genomes of cultured bacterial strains and MAGs (ANIm > 94%; alignment coverage > 10%). (**e**) Comparison of PULs conserved between MAG Bg0018 and Bg0019 with PULs in *P. copri* Bg131 and other *P. copri* MAGs. (**f**) *P. copri* Bg131 PULs. (**g**) Global distribution of *P. copri* Bg131, *P. copri* MAGs, and *P. stercorea* CAZymes in different GH, PL, CE, and CBM families.

**Supplementary Table 2. Diets used in gnotobiotic mouse studies**. (**a**) Ingredients in each diet module. (**b**) Representation of modules in the weaning diet supplemented with MDCF-2. (**c**) Nutritional analysis of diets.

**Supplementary Table 3. Absolute abundances of bacterial strains in dam-pup dyads colonized with cultured bacterial consortia in the experiment described in** Fig. 1. (**a**) Sample metadata. (**b**) Temporal changes in the absolute abundances of strains in fecal samples collected from dams [log_10_(genome equivalents/milligram feces)]. (**c**) Temporal changes in the absolute abundances of bacterial strains in fecal samples collected from progeny of dams [log_10_(genome equivalents/milligram feces)]. (**d**) Absolute abundance of strains [log_10_(genome equivalents/milligram sample)] in ileal and cecal contents collected from P53 progeny of dams.

**Supplementary Table 4. Bacterial strains used in the gnotobiotic mouse experiments shown in Extended Data Fig. 1.**

**Supplementary Table 5. Absolute abundances of bacterial strains in dam-pup dyads colonized with cultured bacterial consortia in the experiment described in Extended Data Fig. 1. (a)** Sample metadata. (**b**) Absolute abundance of strains [log_10_(genome equivalents per gram sample)] in cecal contents collected from progeny of dams at euthanasia.

**Supplementary Table 6. UHPLC-QqQ-MS analysis of the levels of glycosidic linkages and monosaccharides in cecal glycans present in mice from the various treatment arms of the experiment described in** Fig. 1. (**a**) Glycosidic linkage composition (peak area in arbitrary units). (**b**) Monosaccharides (μg/mg lyophilized cecal sample).

**Supplementary Table 7. Cecal microbial RNA-Seq dataset generated from the gnotobiotic mouse experiment described in** Fig. 1. (**a**) Sample metadata. (**b**) Gene Set Enrichment Analysis (GSEA) of predicted PULs from *P. copri* Bg131 and *P. stercorea*. (**c**) Differentially expressed genes (q-value < 0.1) with annotations in mcSEED pathways for carbohydrate utilization, amino acid, and vitamin biosynthesis. (**d**) Gene Set Enrichment Analysis (GSEA) of mcSEED pathways.

**Supplementary Table 8. Quantification of jejunal villus height and crypt depth in gnotobiotic mice in the experiment described in Fig. 1**.

**Supplementary Table 9. snRNA-Seq dataset generated from the jejunums of gnotobiotic mice in the experiment described in** Fig. 1. (**a**) Sample metadata. (**b**) Differentially expressed genes from pseudobulk analysis (q-value < 0.05). (**c**) Proportional representation of cell clusters identified from snRNA-seq dataset.

**Supplementary Table 10. Absolute abundances of bacterial strains in dam-pup dyads colonized with cultured bacterial consortia in the experiment described in Extended Data Fig. 7.** (**a**) Sample metadata. (**b**) Absolute abundance of strains [log_10_(genome equivalents per gram)] in cecal contents collected from P53 offspring of the dams.

**Supplementary Table 11. Targeted LC-MS analysis of levels of acylcarnitines, amino acids and biogenic amines in gnotobiotic mice in the experiment described in Extended Data Fig. 7.** (**a**) Amino acids/other biogenic amines (µM). (**b**) Acylcarnitines (nM). (**c**) Plasma non-esterified fatty acids (NEFA; mmol/L).

**Supplementary Table 12. Comparisons of genomes of *P. copri* isolates BgD5_2 and BgF5_2 to *P. copri* MAGs Bg0018 and Bg0019.** (**a**) Quality metrics for genomes assembled from *P. copri* isolates BgD5_2 and BgF5_2. (**b**) Comparison of mcSEED metabolic pathway annotations between the genomes of P. copri isolates and MAGs Bg0018 and Bg0019. (**c**) Comparison of PUL content between the genomes of *P. copri* isolates and MAGs Bg0018 and Bg0019. (**d**) Comparison of *P. copri* BgD5_2 and BgF5_2 genome identity and uniqueness.

**Supplementary Table 13. Absolute abundances of bacterial strains in gnotobiotic mice colonized with cultured bacterial consortia in the experiment described in** Fig. 4. (**a**) Sample metadata. (**b**) Absolute abundance of bacterial strains [log_10_(genome equivalents per gram)] in cecal contents collected from P53 offspring of the dams.

**Supplementary Table 14. Targeted LC-MS analysis of levels of acylcarnitines, amino acids and biogenic amines in gnotobiotic mice in the experiment described in** Fig. 4. (**a**) Amino acid and biogenic amine results for jejunum (µM). (**b**) Acylcarnitine results for jejunum, ileum, colon, and liver (nM).

**Supplementary Table 15. Cecal microbial RNA-Seq dataset generated from gnotobiotic mice in the experiment described in** Fig. 4. (**a**) Sample metadata. (**b**) Gene Set Enrichment Analysis (GSEA) of predicted PULs from BgD5_2 and BgF5_2.

**Supplementary Table 16. UHPLC-QqQ-MS analysis of levels of glycosidic linkages and monosaccharides in cecal glycans recovered from gnotobiotic mice in the experiment described in** Fig. 4. (**a**) Glycosidic linkage composition (peak area in arbitrary units). (**b**) Monosaccharides (μg/mg lyophilized cecal contents).

**Supplementary Table 17. Targeted UHPLC-QqQ-MS analysis of amino acids and B vitamins in cecal contents collected from gnotobiotic mice in the experiment described in** Fig. 4**. (a)** Amino acids (µg/g of cecal contents). (**b**) B vitamins (ng/g of cecal contents).

